# Model selection in ADMIXTURE can be inconsistent: proof of the *K* = 2 phenomenon

**DOI:** 10.64898/2026.02.27.708651

**Authors:** Dat Do, Jonathan Terhorst

## Abstract

STRUCTURE and ADMIXTURE are two popular methods for detecting population structure in genetic data. They model observed genotypes as mixtures of latent ancestral populations, and the inferred admixture proportions can be used to visualize and summarize population structure. A key parameter in these models is the number of ancestral populations, *K*. Selecting *K* is a challenging problem. Perhaps the most widely used method is Evanno’s Δ*K*, which selects *K* based on the second-order change in log-likelihood as *K* increases. However, practitioners have often noted that Δ*K* often favors overly small *K*, frequently returning *K* = 2 even when more meaningful substructure is present. In this paper, we provide a theoretical explanation for this phenomenon: we prove that, under certain conditions, the Δ*K* method can be inconsistent, meaning that it can fail to identify the true number of populations even with infinite data.

## 1 Introduction

A common first step when exploring genetic data is to group or *cluster* sampled genomes into a smaller number of interpretable units. Two workhorse methods for clustering genetic data predominate. Principal components analysis (PCA) provides a fast, model-free representation of major axes of genetic variation [Patterson et al., 2006, Price et al., 2006]. Alternatively, model-based methods such as STRUCTURE [Pritchard et al., 2000] and ADMIXTURE [Alexander et al., 2009] posit a fixed number *K* of ancestral populations, each with specific allele frequencies, and represent sampled genomes as a mixture over these populations.

An appeal of model-based approaches is interpretability: the inferred clusters represent hypothesized ancestral populations, and individual mixing proportions correspond to ancestry fractions. But this interpretation depends on specifying the number of populations *K*, which is usually non-obvious. Setting *K* too small (*underfitting*) can smooth out real structure, while setting *K* too large (*overfitting*) can turn noise into apparent structure. Unlike PCA, where dimensionality can be explored *post hoc*, model-based approaches require this choice in advance, and different criteria can yield different conclusions.

Despite the widespread use of STRUCTURE and ADMIXTURE, the question of how best to select *K* remains unsettled [Novembre, 2016]. Perhaps the most widely used approach is Evanno’s Δ*K* [Evanno et al., 2005], which detects an “elbow” in a likelihood summary as *K* increases. However, practitioners have noted that Δ*K* often favors overly small *K*, especially *K* = 2. Indeed, a recent survey of 1,264 studies that utilized STRUCTURE found that use of Δ*K* increased over time and was associated with a substantially higher rate of reporting *K* = 2 [Janes et al., 2017], with potentially serious implications for species conservation and management. A follow-up simulation study supported the tendency of Δ*K* to select *K* = 2, and found that the magnitude of Δ*K* is related to the level of divergence between populations [Cullingham et al., 2020]. This phenomenon is also noted in the seminal paper [Evanno et al., 2005].

Although the tendency of Δ*K* to underfit has been documented empirically, it lacks a rigorous mathematical explanation. Here we provide one, by proving that Δ*K* can be *inconsistent*: in some regimes, even with infinite data, it selects *K* = 2 despite the true value being larger. We give an explicit sufficient condition for this to occur in terms of the probabilistic divergence between population allele frequency distributions, and show how this condition can be satisfied under a realistic population genetic model when *F*_*ST*_ is sufficiently low. To the best of our knowledge, this is the first theoretical explanation for the *K* = 2 phenomenon.

## 2 Results

In this study, we focus on maximum likelihood estimation (MLE) of the STRUCTURE model, i.e. the ADMIXTURE method [Alexander et al., 2009]. We quickly review the model in order to be able to state our results.

ADMIXTURE assumes genotype matrix 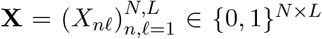 of *N* individuals at *L* SNPs is generated by admixing between *K*_0_ “true” populations^1^. Each population *k* (*k* ∈ [*K*_0_]) is represented by its allele frequency vector 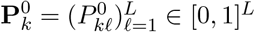. Each individual *n* ∈ {1, …, *N*} is a mixture of those populations, with proportion 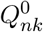 belonging to population *k*, where 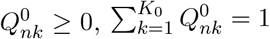. Hence, it models the genotype data as

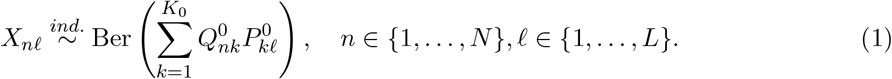

The parameters 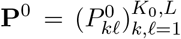 and 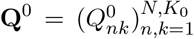 and the number of populations *K*_0_ are unknown and have to be estimated from data.

### 2.1 Model selection using second-order changes of log-likelihood functions

Now we present an Evanno-style “elbow” criterion for selecting *K* adapted to the MLE framework. For a given *K*, the average log-likelihood of the model with parameters (**Q, P**) is

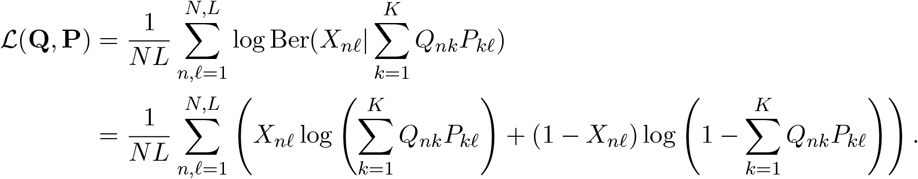

For every *K* ranging from 1 to some pre-specified upper bound 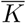, we seek the MLE

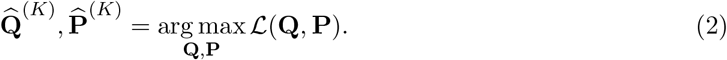

Let 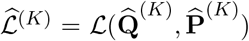. For 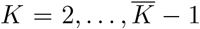, we examine the second-order change of these log-likelihood values as

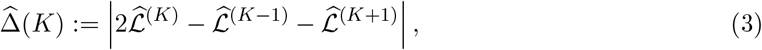

and decide the most plausible value of *K* as

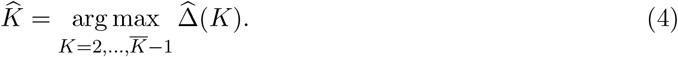

Note that our definition of 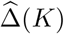 differs from the original Δ*K* in [Evanno et al., 2005], which divides by a standard-deviation term estimated across runs. We analyze the un-normalized form because the normalized statistic is a ratio of dependent random variables, hence difficult to analyze theoretically. Also, whereas this normalization was originally intended to prevent *overfitting* (over-fitted log-likelihoods have higher variance), at least in the scenarios considered here, the Δ*K* method *under*fits, so this normalization only exacerbates the problem.

### 2.2 Inconsistency of model selection

We focus on the case of true *K*_0_ = 3 populations and denote the true parameters by **Q**^0^ ∈ [0, 1]^*N×*3^ and **P**^0^ ∈ [0, 1]^3*×L*^. We state informal asymptotic versions of the two main theorems here, and defer full non-asymptotic statements and proofs to the SI Appendix. Our results make the following assumptions about the data generating process.

#### Assumption 1

(Boundedness of the allele frequency **P**^0^). There exists a positive constant *c* so that true parameters 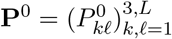 satisfies

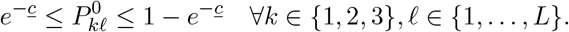

Assumption 1 ensures that the allele frequencies are bounded away from 0 and 1. This is necessary to keep the log-likelihood terms from diverging near the boundaries of the parameter space. It is not overly restrictive, since it is common practice to QC-filter rare variants (low minor allele frequency), which are more error-prone and often less informative for PCA or ADMIXTURE analyses.

#### Assumption 2

(Data generating assumption of **Q**^0^). *N* samples are partitioned into three groups 𝒩_1_ ∪ 𝒩_2_ ∪ 𝒩_3_ so that 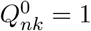 for every *n* ∈ 𝒩_*k*_ and *k* ∈ [3]. We also assume |𝒩_1_| = |𝒩_2_| = |𝒩_3_| = *N/*3.

Assumption 2 places each individual at a vertex of the simplex (i.e., 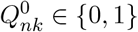). It is made for mathematical convenience, but it is also favorable for identifying population structure: every individual is purely a member of a single population, which maximizes the signal of population structure in the data.

#### 2.2.1 Measures of population divergence

Our first theorem is stated in terms of the Kullback-Leibler (KL) divergence between allele frequencies across populations. KL divergence emerges naturally when studying the asymptotics of likelihood-based inference methods. In the sequel, we offer a second result phrased in terms of population genetic divergences (i.e., *F*_*ST*_) that may be more familiar to practitioners.

For two populations with allele frequencies **P**_1_ and **P**_2_ ∈ [0, 1]^*L*^, let

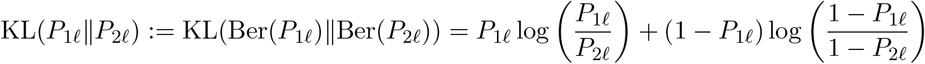

denote the Kullback-Leibler divergence between allele frequencies of SNP *f* for all *f* ∈ [*L*], and let

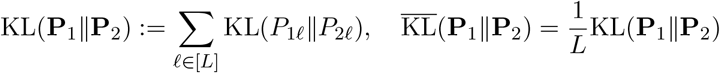

denote their sum and average over *f*, respectively. We suppose a tree-like structure for three populations so that 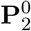 and 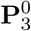 are closer to each other compared to 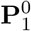, and this is quantified via

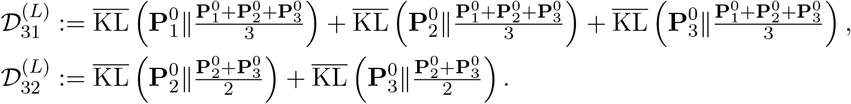

The population-level analogs of 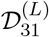 and 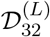 are obtained by averaging over the locus distribution (equivalently, by taking *L* → ∞):

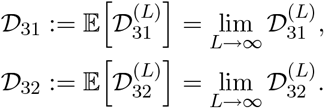

#### 2.2.2 First main result

Our first main result shows that Evanno’s second-order criterion can underfit in the idealized three-population setting described above.

##### Theorem 1.

*Suppose that*

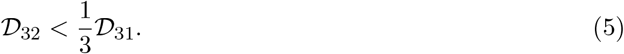

*Then, as N, L* →∞, *the second-order method* (4) *selects* 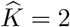 *with probability tending to 1 (despite the true K*_0_ = 3*), i*.*e*., *it is inconsistent*.

𝒟_32_ is the average information loss from merging populations 2 and 3, while 𝒟_31_ captures overall three-way heterogeneity. Condition (5) therefore says that merging 2 and 3 is cheap relative to total dispersion, so the elbow criterion can prefer 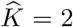 even asymptotically. The SI Appendix formalizes this by relating fitted log-likelihood gaps to their population counterparts with error *c*_*NL*_ → 0 as *N, L* → ∞.

Note that Theorem 1 conditions on **P**^0^ and **Q**^0^, and does not assume any particular mechanism for producing them. Thus any process that generates parameters satisfying the divergence inequality (5) can lead to inconsistency of Δ*K*. In the next section we provide an example of such a process based on a realistic population-genetic model.

#### 2.2.3 Second main result

Our next theorem shows how the divergence inequality (5) can be satisfied when the data are generated under a hierarchical population model with realistic covariance [Balding and Nichols, 1995]. At each locus *ℓ*, let 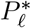 denote the root allele frequency. Conditional on 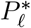, we draw an internal node 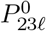 and three leaves 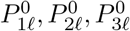 via

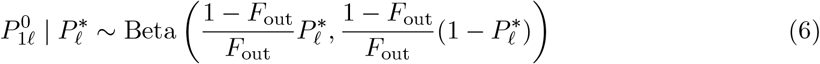

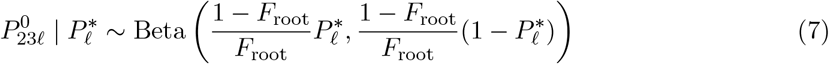

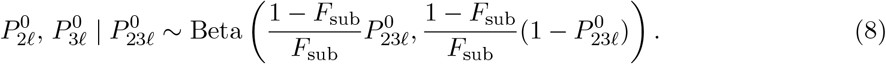

Here *F*_root_, *F*_sub_, *F*_out_ are drift parameters along the edges 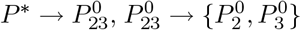, and 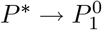, respectively (see Figure 1). To match the conditional variances of the leaves, so that 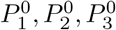 are equally dispersed around *P* ^∗^, we set

**Figure 1.**
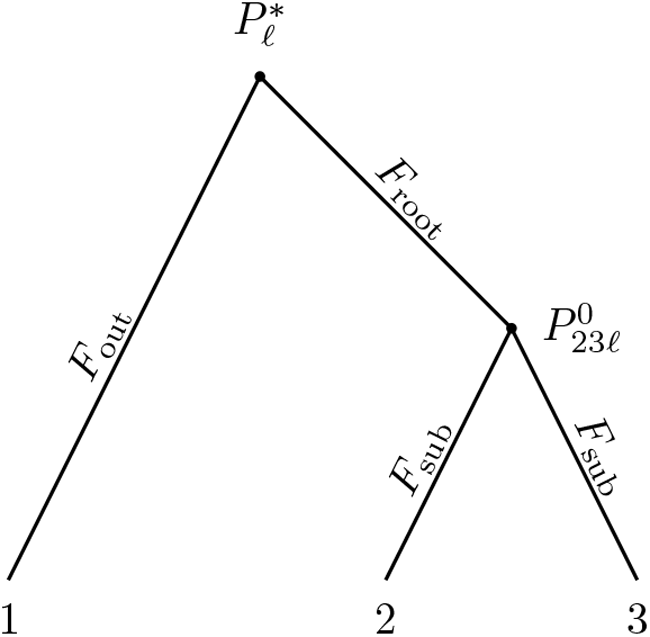
Nested Balding–Nichols model with drift parameters on branches.

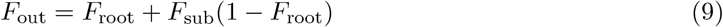

##### Theorem 2.

*Suppose the population allele frequencies are generated according to the nested Balding-Nichols model* (6)*–*(9), *and that the ancestral allele frequencies* 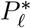 *are bounded away from 0 and 1. If F*_*root*_ *and F*_*sub*_ *are sufficiently small and*

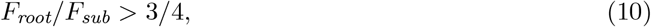

*the* Δ*K method selects* 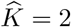 *with probability tending to 1 as N, L* → ∞.

The proof of Theorem 2 relies on a “small drift” argument by expanding the extant allele frequencies 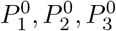 around *P* ^∗^ to first-order in *F*_root_ and *F*_sub_.

We performed numerical simulations to check the predicted threshold (10). We considered *F*_sub_ ∈ {0.01, 0.05, 0.2} and 33 values of *F*_root_ for each *F*_sub_ (99 settings total), including a dense set of ratios near the theoretical boundary *F*_root_*/F*_sub_ = 3*/*4. This range of *F*_*ST*_ covers values typically seen in human populations [Bhatia et al., 2013]. As *F*_root_ → 0 the demography approaches a star topology, while for larger values, the relationship becomes increasingly hierarchical. For each parameter setting, we simulated *L* = 2000 SNPs and *N* = 150 diploid individuals (50 from each of the three populations). For each simulated data set, we fit ADMIXTURE for *K* = 1, …, 5 and retained the maximum log-likelihood at each *K* across multiple random restarts. We then compute 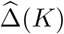 for each *K* and select 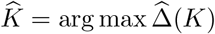.

Figure 2A displays 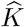 against log_10_(*F*_root_*/F*_sub_), and Figure 2B shows the same outcomes in the (*F*_sub_, *F*_root_) plane. The simulation results in Figure 2 show a clear transition near the predicted boundary (10). It is interesting to note that when the distance *F*_sub_ between 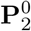 and 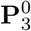 is fixed and *F*_root_ is small, 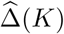 can correctly choose 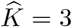. But when *F*_root_ increases, 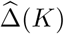 favors merging 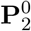 and 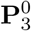 into one population.

**Figure 2.**
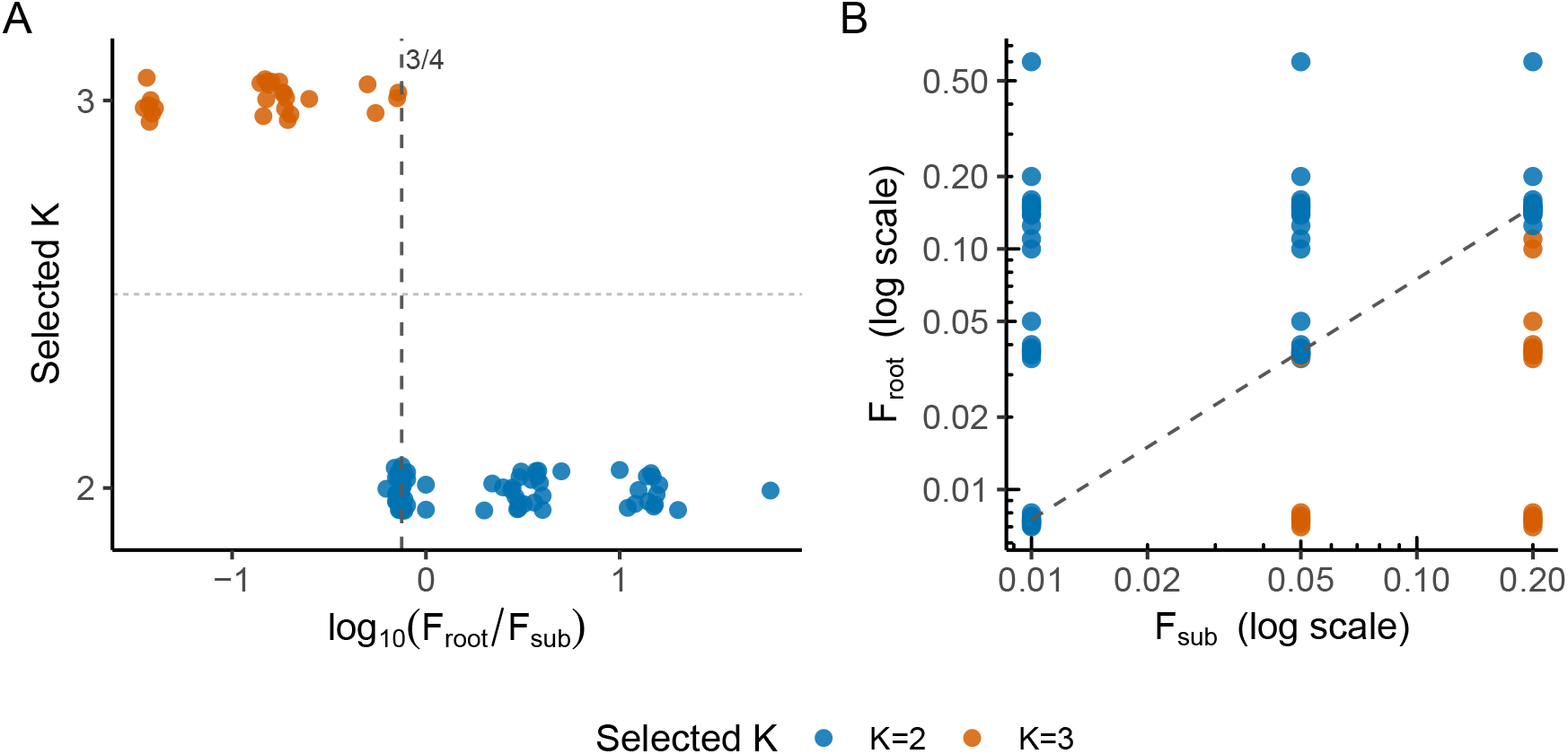
Threshold phenomenon predicted by Theorem 2. **(A)** Across simulated three-population settings under the nested Balding–Nichols model, the 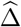 method selects 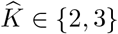 as a function of log_10_(*F*_root_*/F*_sub_). **(B)** The same settings shown in the (*F*_sub_, *F*_root_) plane (log scale). The dashed line in each plot marks the theoretical phase boundary *F*_root_*/F*_sub_ = 3*/*4.

## 3 Discussion

We provide a theoretical explanation for the widely observed tendency of Evanno’s Δ*K* to underfit in ADMIXTURE analyses. In particular, we identify regimes where Δ*K* is inconsistent, selecting 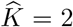 even when the true number of populations is 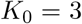, and show how this could arise in a realistic three-population model with *F*_*ST*_ values similar to those observed in modern humans.

Our results do not imply that Δ*K* always fails. They identify a specific failure mode, most relevant when populations are closely related. In practice, Δ*K* should be interpreted alongside other selection criteria and biological context, and results across a range of *K* should be reported rather than relying on a single selected value, as advocated by prior work [Pritchard et al., 2000, Evanno et al., 2005, Janes et al., 2017].

Here we focused on maximum likelihood estimation and the Δ*K* method. In practice, other model selection techniques and inference methods are often used to fit the ADMIXTURE model. Nevertheless, many if not all of them rely on comparing log-likelihoods across different *K* to perform model selection. It is therefore plausible to conjecture that they are vulnerable to the same underfitting phenomenon when populations are closely related. We leave a rigorous analysis of these other methods to future work.

## Acknowledgments

Dat Do is grateful to Prof. Long Nguyen for stimulating discussions on empirical process theory. This research was supported in part by the National Institute of General Medical Sciences of the NIH under award number R35GM151145. The content is solely the responsibility of the authors and does not necessarily represent the official views of the NIH.

## Footnotes

1 To simplify the argument, we present the haploid admixture model, but the results can be easily generalized to polyploid organisms with similar results.

## Notes

### Competing Interest Statement

The authors have declared no competing interest.

