## Supporting Information for "Model selection in ADMIXTURE can be inconsistent: proof of the *K* = 2 phenomenon"

Here we provide formal versions of Theorems 1 and 2, as well as some supplementary results and discussion. We recall for reference the following assumptions from the main text:

**Assumption 1** (Data generating assumption of $\mathbf{Q}^0$)**.** $N$ samples are partitioned into three groups $\mathcal{N}_1 \cup \mathcal{N}_2 \cup \mathcal{N}_3$ so that $Q^0_{nk} = 1$ for every $n \in \mathcal{N}_k$ and $k \in [3]$. We also assume $|\mathcal{N}_1| = |\mathcal{N}_2| = |\mathcal{N}_3| = N/3$.

**Assumption 2** (Boundedness of the allele frequency $\mathbf{P}^0$)**.** There exists a positive constant $\underline{c}$ so that true parameters $\mathbf{P}^0 = (P^0_{k\ell})^{3,L}_{k,\ell=1}$ satisfies

$$e^{-\underline{c}} \leq P^0_{k\ell} \leq 1 - e^{-\underline{c}} \quad \forall k \in [3], \ell \in [L].$$

For a natural number $n$, we use $[n]$ to denote $\{1, \dots, n\}$.

#### 1 Theorem 1

We now state our main non-asymptotic result about the inconsistency of the $\widehat{\Delta}(K)$ method. All probability statements are with respect to the randomness of the sample $\mathbf{X} \in \{0,1\}^{N \times L}$ generated according to the true admixture model's parameters $\mathbf{Q}^0$ and $\mathbf{P}^0$.

**Theorem 1.** *Given Assumption 1 and Assumption 2, and assume further that the true allele frequency $\mathbf{P}^0$ satisfies:*

$$\overline{KL}\left(\boldsymbol{P}^0_2 \middle\| \frac{\boldsymbol{P}^0_2 + \boldsymbol{P}^0_3}{2}\right) + \overline{KL}\left(\boldsymbol{P}^0_3 \middle\| \frac{\boldsymbol{P}^0_2 + \boldsymbol{P}^0_3}{2}\right) + \epsilon_{NL} < \frac{1}{3}\sum_{j=1}^{3} \overline{KL}\left(\boldsymbol{P}^0_j \middle\| \frac{\boldsymbol{P}^0_1 + \boldsymbol{P}^0_2 + \boldsymbol{P}^0_3}{3}\right), \tag{1}$$

*where*

$$\epsilon_{NL} = 8\frac{C\overline{K}(N+L)\log(NL)}{NL} + 4\underline{c}\left(\frac{\log(NL)}{NL}\right)^{1/2} + 2\left(\frac{\log(L)}{L}\right)^{1/2} + 2C_1\left(\frac{\log(N)}{N}\right)^{1/2} \tag{2}$$

*then*

$$\mathbb{P}(\widehat{K} = 2) \geq 1 - 3(NL)^{-c_0} - 4L^{-4e^2}, \tag{3}$$

*for all sufficiently large $N$ and $L$, where $C, C_1$ and $c_0$ are universal positive constants.*

**Remark 1.** *Note that the left-hand side (LHS) of the core assumption (1) only depends on $\boldsymbol{P}^0_2$ and $\boldsymbol{P}^0_3$, and we can choose $\boldsymbol{P}^0_1$ sufficiently far away from them for (1) to hold. As $N, L \to \infty$, the $\epsilon_{NL}$ term in the LHS also tends to 0, and the probability that $\widehat{K} = 2$ tends to 1. Because the true $K_0 = 3$, it indicates that $\widehat{\Delta}(K)$ is under-fitted (therefore inconsistent) as the number of observations tends to infinity.*

We summarize this discussion in a corollary, which is presented as the informal Theorem 1 in the main text.

**Corollary 1.** *Given Assumption 1 and Assumption 2, and assume that there is a positive constant $C_g$ such that*

$$\overline{KL}\left(\boldsymbol{P}_2^0 \middle\| \frac{\boldsymbol{P}_2^0+\boldsymbol{P}_3^0}{2}\right) + \overline{KL}\left(\boldsymbol{P}_3^0 \middle\| \frac{\boldsymbol{P}_2^0+\boldsymbol{P}_3^0}{2}\right) + C_g < \frac{1}{3}\sum_{j=1}^{3} \overline{KL}\left(\boldsymbol{P}_j^0 \middle\| \frac{\boldsymbol{P}_1^0+\boldsymbol{P}_2^0+\boldsymbol{P}_3^0}{3}\right), \tag{4}$$

*for every $L$, then*

$$\mathbb{P}(\widehat{K} < K_0) \to 1, \tag{5}$$

*as $N$ and $L$ tend to infinity, i.e., $\widehat{\Delta}(K)$ method is inconsistent.*

**Remark 2.** *The term on the left-hand side of the core inequality (1)*

$$\mathcal{D}_{32}^{(L)} := \left(\overline{KL}\left(\boldsymbol{P}_2^0 \| \tfrac{\boldsymbol{P}_2^0+\boldsymbol{P}_3^0}{2}\right) + \overline{KL}\left(\boldsymbol{P}_3^0 \| \tfrac{\boldsymbol{P}_2^0+\boldsymbol{P}_3^0}{2}\right)\right)$$

*is twice the Jensen-Shannon divergence between $\boldsymbol{P}_2^0$ and $\boldsymbol{P}_3^0$ (average across L SNPs). The right-hand side, which is denoted by $\mathcal{D}_{31}^{(L)}$, is the average Jensen-Shannon divergence between $\boldsymbol{P}_1^0$, $\boldsymbol{P}_2^0$, and $\boldsymbol{P}_3^0$. An important step in the proof of Theorem 1 involves bounding the difference in maximal log-likelihoods at different $K$ in admixture models by $\mathcal{D}_{32}^{(L)}$ and $\mathcal{D}_{31}^{(L)}$, i.e.,*

$$\mathcal{L}^{(3)} - \mathcal{L}^{(2)} \leq \mathcal{D}_{32}^{(L)}, \quad \text{and} \quad \mathcal{L}^{(3)} - \mathcal{L}^{(1)} \geq \mathcal{D}_{31}^{(L)}, \tag{6}$$

*with a probability tending to 1. KL divergence and Jensen-Shannon divergence naturally arise in likelihood-based inference, such as the MLE procedure we are considering. Later in Section 2 and 3, we will relate them to the more popular notions of distance in population genetics, such as $F_2, F_3$, and $F_{ST}$.*

#### 1.1 Proof of the theorem

Figure 1 shows an overview of the proof of Theorem 1. It uses three supporting lemmas (Lemma 1, 2, and 3) to control the gap between the average log-likelihood of the model under different $K$. They will be presented and proved after the proof of the main theorem. Recall that for a set of admixture proportions $\mathbf{Q} \in [0,1]^{N\times K}$ and allele frequencies $\mathbf{P} \in [0,1]^{K\times L}$, the average log-likelihood function is denoted as:

$$\begin{aligned}\mathcal{L}(\mathbf{Q},\mathbf{P}) &= \frac{1}{NL}\sum_{n,\ell=1}^{N,L} \log \mathrm{Ber}(X_{n\ell}|\sum_{k=1}^{K} Q_{nk}P_{k\ell}) \\ &= \frac{1}{NL}\sum_{n,\ell=1}^{N,L}\left(X_{n\ell}\log\left(\sum_{k=1}^{K} Q_{nk}P_{k\ell}\right) + (1-X_{n\ell})\log\left(1-\sum_{k=1}^{K} Q_{nk}P_{k\ell}\right)\right).\end{aligned}$$

The data $\mathbf{X} \in \{0,1\}^{N\times L}$ is generated from true parameters $\mathbf{Q}^0 \in [0,1]^{N\times 3}$ and $\mathbf{P}^0 \in [0,1]^{3\times L}$ as $X_{n\ell} \sim \mathrm{Ber}(Q_{n1}^0 P_{1\ell}^0 + Q_{n2}^0 P_{2\ell}^0 + Q_{n3}^0 P_{3\ell}^0)$ and $K_0 = 3$. For each $K \in [1, \overline{K}]$, let $\mathbf{Q}^{(K)}$ and $\mathbf{P}^{(K)}$ be the MLE, and the maximal average log-likelihood is denoted as $\widehat{\mathcal{L}}^{(K)} = \mathcal{L}(\mathbf{Q}^{(K)}, \mathbf{P}^{(K)})$.

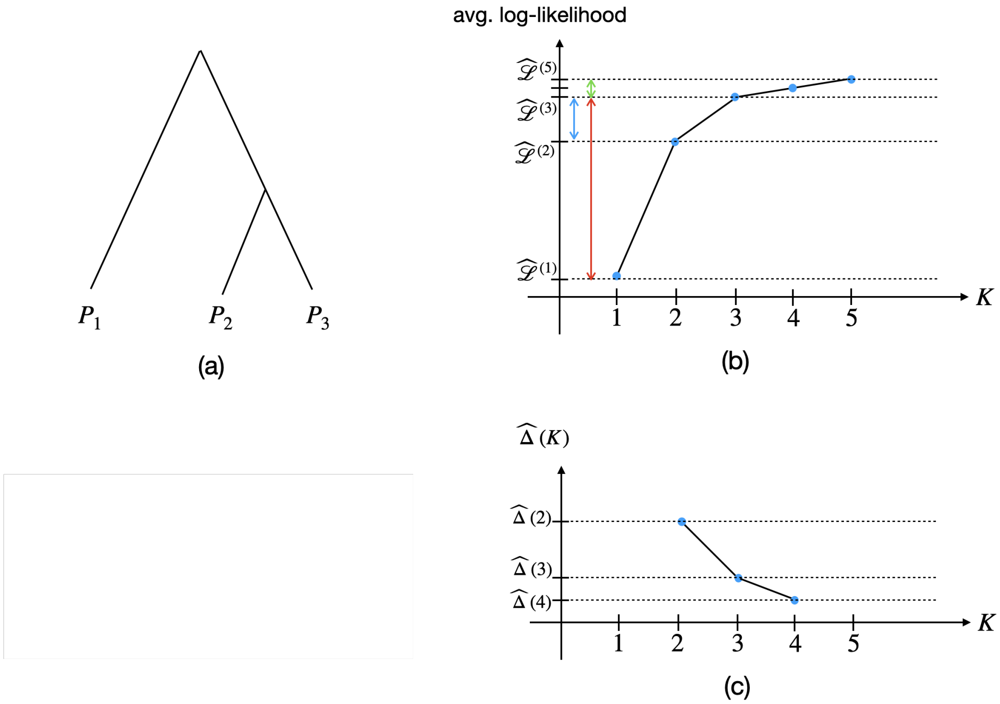


Figure 1: Overview of the theoretical results. **Panel (a):** We consider three populations where $P_2$ and $P_3$ are more closely related compared to $P_1$. **Panel (b):** Proving Theorem 1 requires three steps (supporting lemmas). Lemma 1 shows that the gap of the average log-likelihood from exact-fitted to over-fitted level (the green segment) tends to 0. Lemma 2 bounds the gap between the log-likelihood of the admixture model with $K = 3$ and $K = 2$ (the blue segment) from above by $\mathcal{D}_{32}^{(L)}$. Lemma 3 bounds the gap between the log-likelihood of the admixture model with $K = 3$ and $K = 1$ (the red segment) from below by $\mathcal{D}_{31}^{(L)}$. **Panel (c):** When $\mathcal{D}_{32}^{(L)} < \frac{1}{3}\mathcal{D}_{31}^{(L)}$ then it implies by a simple algebraic argument that $\widehat{\Delta}(2) > \widehat{\Delta}(3)$ as $N, L \to \infty$, which indicates that $\widehat{\Delta}$ is inconsistent (see the proof of Theorem 1).

*Proof of Theorem 1.* Our goal is to show that the second-order change at $K = 2$, i.e., $\widehat{\Delta}(2)$ is the largest over all $\widehat{\Delta}(K)$ (for $K = 2, \ldots, \overline{K} - 1$) with the probability tending to 1 as $N$ and $L \to \infty$. The proof is divided into 3 steps. At a high level, in Step 1, we will show that

$$\widehat{\Delta}(2) > \widehat{\Delta}(3), \tag{7}$$

and in Step 2, we prove

$$\widehat{\Delta}(2) > \widehat{\Delta}(K), \tag{8}$$

for all $K \geq 4$, all with probability tending to 1. Then we combine them to conclude that $\widehat{\Delta}(2)$ is the largest, i.e., the model selection method using $\widehat{\Delta}$ will select $\widehat{K} = 2$ as the number of observations tends to infinity.

**Step 1: Proving inequality (7).**

The inequality $\widehat{\Delta}(2) > \widehat{\Delta}(3)$ can be expressed as

$$|2\widehat{\mathcal{L}}^{(2)} - \widehat{\mathcal{L}}^{(1)} - \widehat{\mathcal{L}}^{(3)}| > |2\widehat{\mathcal{L}}^{(3)} - \widehat{\mathcal{L}}^{(2)} - \widehat{\mathcal{L}}^{(4)}|. \tag{9}$$

We first prove the inequality without the absolute value sign, i.e.,

$$2\widehat{\mathcal{L}}^{(2)} - \widehat{\mathcal{L}}^{(1)} - \widehat{\mathcal{L}}^{(3)} > 2\widehat{\mathcal{L}}^{(3)} - \widehat{\mathcal{L}}^{(2)} - \widehat{\mathcal{L}}^{(4)}. \tag{10}$$

Re-arrange this inequality and note that $\widehat{\mathcal{L}}^{(4)} \geq \widehat{\mathcal{L}}^{(3)}$, it suffices to prove

$$3\widehat{\mathcal{L}}^{(2)} > 2\widehat{\mathcal{L}}^{(3)} + \widehat{\mathcal{L}}^{(1)}, \tag{11}$$

which is equivalent to

$$3\left(\widehat{\mathcal{L}}^{(3)} - \widehat{\mathcal{L}}^{(2)}\right) < \widehat{\mathcal{L}}^{(3)} - \widehat{\mathcal{L}}^{(1)}. \tag{12}$$

By Lemma 2, there exists an event $A$ with $\mathbb{P}(A) \geq 1 - (NL)^{-c} - (NL)^{-c_1}$ so that for all $\mathbf{X} \in A$, it holds that

$$\widehat{\mathcal{L}}^{(3)} - \widehat{\mathcal{L}}^{(2)} \leq \frac{1}{3}\left(\overline{\mathrm{KL}}\left(\mathbf{P}_2^0 \middle\| \frac{\mathbf{P}_2^0 + \mathbf{P}_3^0}{2}\right) + \overline{\mathrm{KL}}\left(\mathbf{P}_3^0 \middle\| \frac{\mathbf{P}_2^0 + \mathbf{P}_3^0}{2}\right)\right) + \delta_{NL} + \underline{c}\left(\frac{\log(NL)}{NL}\right)^{1/2}, \tag{13}$$

where $c$ and $c_1$ are universal constants, and

$$\delta_{NL} := \frac{C\overline{K}(N+L)\log(NL)}{NL},$$

where $C$ is also a universal constant (cf. Lemma 1). Moreover, from Lemma 3, we have another event $B$ with $\mathbb{P}(B) \geq 1 - (NL)^{-c_2} - 4L^{-4e^2}$ such that

$$\widehat{\mathcal{L}}^{(3)} - \widehat{\mathcal{L}}^{(1)} \geq \frac{1}{3}\sum_{k=1}^{3}\overline{\mathrm{KL}}(\mathbf{P}_k^0 \| \tfrac{\mathbf{P}_1^0 + \mathbf{P}_2^0 + \mathbf{P}_3^0}{3}) - \underline{c}\left(\frac{\log(NL)}{NL}\right)^{1/2} - 2\left(\frac{\log(L)}{L}\right)^{1/2} - 2C_1\left(\frac{\log(N)}{N}\right)^{1/2}, \tag{14}$$

where $c_2$ and $C_1$ are universal positive constants. By Bonferroni's bound, we have $\mathbb{P}(A \cap B) \geq \mathbb{P}(A) + \mathbb{P}(B) - 1 \geq 1 - ((NL)^{-c} + (NL)^{-c_1} + (NL)^{-c_2} + 4L^{-4e^2})$. On the event $A \cap B$, both inequalities above hold and by combining with the assumed inequality (19), we have

$$3\left(\widehat{\mathcal{L}}^{(3)} - \widehat{\mathcal{L}}^{(2)}\right) \leq \overline{\mathrm{KL}}\left(\mathbf{P}_2^0 \middle\| \frac{\mathbf{P}_2^0 + \mathbf{P}_3^0}{2}\right) + \overline{\mathrm{KL}}\left(\mathbf{P}_3^0 \middle\| \frac{\mathbf{P}_2^0 + \mathbf{P}_3^0}{2}\right) + 3\delta_{NL} + 3\underline{c}\left(\frac{\log(NL)}{NL}\right)^{1/2}$$

$$
\begin{aligned}
&< \frac{1}{3}\sum_{k=1}^{3}\overline{\mathrm{KL}}(\mathbf{P}_k^0\|\tfrac{\mathbf{P}_1^0+\mathbf{P}_2^0+\mathbf{P}_3^0}{3}) - \underline{c}\left(\frac{\log(NL)}{NL}\right)^{1/2} - 2\left(\frac{\log(L)}{L}\right)^{1/2} - 2C_1\left(\frac{\log(N)}{N}\right)^{1/2} \\
&\le \widehat{\mathcal{L}}^{(3)} - \widehat{\mathcal{L}}^{(1)}.
\end{aligned}
$$

Hence, (12) is correct, which implies that (10) holds in the event $A \cap B$. Now we proceed to show the inequality with absolute value signs (9). Let $\alpha = 2\widehat{\mathcal{L}}^{(2)} - \widehat{\mathcal{L}}^{(1)} - \widehat{\mathcal{L}}^{(3)}$ and $\beta = 2\widehat{\mathcal{L}}^{(3)} - \widehat{\mathcal{L}}^{(2)} - \widehat{\mathcal{L}}^{(4)}$,

$$
\begin{aligned}
\alpha + 2\beta &= 3\widehat{\mathcal{L}}^{(3)} - \widehat{\mathcal{L}}^{(1)} - 2\widehat{\mathcal{L}}^{(4)} \\
&\ge \widehat{\mathcal{L}}^{(3)} - \widehat{\mathcal{L}}^{(1)} - 2\delta_{NL} \\
&\ge \frac{1}{3}\sum_{k=1}^{3}\overline{\mathrm{KL}}(\mathbf{P}_k^0\|\tfrac{\mathbf{P}_1^0+\mathbf{P}_2^0+\mathbf{P}_3^0}{3}) - \underline{c}\left(\tfrac{\log(NL)}{NL}\right)^{1/2} - 2\left(\tfrac{\log(L)}{L}\right)^{1/2} - 2C_1\left(\tfrac{\log(N)}{N}\right)^{1/2} - 2\delta_{NL} \\
&> \overline{\mathrm{KL}}\left(\mathbf{P}_2^0\middle\|\frac{\mathbf{P}_2^0+\mathbf{P}_3^0}{2}\right) + \overline{\mathrm{KL}}\left(\mathbf{P}_3^0\middle\|\frac{\mathbf{P}_2^0+\mathbf{P}_3^0}{2}\right) + 6\delta_{NL} > 0, \qquad (15)
\end{aligned}
$$

where we the inequality $\widehat{\mathcal{L}}^{(3)} \ge \widehat{\mathcal{L}}^{(4)} - \delta_{NL}$ in the event $A$ (Lemma 1 and Lemma 2). Hence, we have shown that $\alpha > -2\beta$ and $\alpha > \beta$, which implies $\alpha > 0$. Furthermore, by noting that $|\beta| = \beta$ if $\beta \ge 0$ and $|\beta| = -\beta$ if $\beta < 0$, those inequalities imply

$$\widehat{\Delta}(2) = |\alpha| = \alpha > |\beta| = \widehat{\Delta}(3).$$

**Step 2. Proving inequality (8).**

For all $K \ge 4$, thanks to Lemma 1 and Lemma 2, we have for all $\mathbf{X} \in A$ that

$$\widehat{\Delta}(K) = 2\widehat{\mathcal{L}}^{(K)} - \widehat{\mathcal{L}}^{(K+1)} - \widehat{\mathcal{L}}^{(K-1)} \ge 2\mathcal{L}(\mathbf{Q}^0, \mathbf{P}^0) - 2\mathcal{L}(\mathbf{Q}^0, \mathbf{P}^0) - \tfrac{2CK(N+L)\log(NL)}{NL} \ge -2\delta_{NL}, \quad (16)$$

and

$$\widehat{\Delta}(K) = 2\widehat{\mathcal{L}}^{(K)} - \widehat{\mathcal{L}}^{(K+1)} - \widehat{\mathcal{L}}^{(K-1)} \le 2\mathcal{L}(\mathbf{Q}^0, \mathbf{P}^0) + \tfrac{2CK(N+L)\log(NL)}{NL} - 2\mathcal{L}(\mathbf{Q}^0, \mathbf{P}^0) \le 2\delta_{NL}, \quad (17)$$

so $|\widehat{\Delta}(K)| \le 2\delta_{NL}$ for all $K > 3$. Moreover, in Step 1, we proved that in the event $A \cap B$, the following inequality holds:

$$\widehat{\Delta}(2) \ge \frac{1}{3}(\widehat{\Delta}(2) + 2\widehat{\Delta}(3)) > 2\delta_{NL}. \qquad (18)$$

Hence,

$$\widehat{\Delta}(2) > \widehat{\Delta}(K) \quad \forall K = 4, \dots, \overline{K} - 1.$$

Combining with the result in Step 1, we have $\widehat{\Delta}(2) = \max\{\widehat{\Delta}(K) : K = 2, \dots, \overline{K} - 1\}$ with probability at least $1 - ((NL)^{-c} + (NL)^{-c_1} + (NL)^{-c_2} + 4L^{-4e^2})$. Let $c_0 = \min\{c, c_1, c_2\}$ we have the conclusion of this theorem. □

#### 1.2 Supporting lemmas

Now we collect and prove all the supporting lemmas of Theorem 1. We first recall a concentration bound for the log-likelihood of exact- and over-fitted MLE that will help us derive the limit of $\widehat{\mathcal{L}}^{(K)}$ for all $K \ge 3$ when $N$ and $L$ get large.

**Lemma 1.** *There exist universal constant $c$ and $C$ such that for all $K \in [3, \overline{K}]$, we have*

$$\mathcal{L}(\boldsymbol{Q}^0, \boldsymbol{P}^0) \leq \widehat{\mathcal{L}}^{(K)} \leq \mathcal{L}(\boldsymbol{Q}^0, \boldsymbol{P}^0) + \frac{CK(N+L)\log(NL)}{NL} \tag{19}$$

*with probability at least $1 - (NL)^{-c}$.*

Let $\delta_{NL} := \dfrac{C\overline{K}(N+L)\log(NL)}{NL}$, we have $\delta_{NL} \to 0$ as $N$ and $L$ get large. Hence, the average log-likelihood of the exact and over-fitted MLE all converge to the average log-likelihood of the true parameters as $N, L \to \infty$. Proving this lemma requires using Empirical Process theory from [van de Geer, 2000] and we deferred the long proof until Section 1.3 for the flow of reading.

Next, the proof of Theorem 1 requires an upper bound $\widehat{\mathcal{L}}^{(3)} - \widehat{\mathcal{L}}^{(2)}$ and a lower bound of $\widehat{\mathcal{L}}^{(3)} - \widehat{\mathcal{L}}^{(1)}$, which are shown in Lemma 2 and Lemma 3.

**Lemma 2.** *Assume Assumption 1 and Assumption 2. Let $\widehat{\mathcal{L}}^{(2)}$ and $\widehat{\mathcal{L}}^{(3)}$ be the maximal average log-likelihood at $K = 2$ and $K = 3$, respectively. Then,*

$$\widehat{\mathcal{L}}^{(3)} - \widehat{\mathcal{L}}^{(2)} \leq \frac{1}{3}\left(\overline{KL}\left(\boldsymbol{P}_2^0 \middle\| \frac{\boldsymbol{P}_2^0 + \boldsymbol{P}_3^0}{2}\right) + \overline{KL}\left(\boldsymbol{P}_3^0 \middle\| \frac{\boldsymbol{P}_2^0 + \boldsymbol{P}_3^0}{2}\right)\right) + \delta_{NL} + \underline{c}\left(\frac{\log(NL)}{NL}\right)^{1/2}. \tag{20}$$

*with a probability exceeding $1 - (NL)^{-c} - (NL)^{-c_1}$, where $c$ and $c_1$ are universal positive constants.*

*Proof.* Due to the concentration inequality in Lemma 1, with probability at least $1 - (NL)^{-c}$ it holds that

$$\begin{aligned}\widehat{\mathcal{L}}^{(K)} &\leq \mathcal{L}(\mathbf{Q}^0, \mathbf{P}^0) + \delta_{NL}, \\ &= \frac{1}{NL}\sum_{\ell=1}^{L}\sum_{k=1}^{3}\sum_{n \in \mathcal{N}_k} X_{n\ell}\log(P_{k\ell}^0) + (1 - X_{n\ell})\log(1 - P_{k\ell}^0) + \delta_{NL},\end{aligned} \tag{21}$$

for all $K = 3, \ldots, \overline{K}$, where the equality is because true admixing proportion $\mathbf{Q}^0$ is assumed to be generated from three populations as in Assumption 1. Now, the fact that $\widehat{\mathcal{L}}^{(2)}$ is the maximal log-likelihood for $K = 2$ implies

$$\begin{aligned}\widehat{\mathcal{L}}^{(2)} \geq \mathcal{L}(\tilde{\mathbf{Q}}^{(2)}, \tilde{\mathbf{P}}^{(2)}) =& \frac{1}{NL}\sum_{\ell=1}^{L}\Bigg[\sum_{n \in \mathcal{N}_1} X_{n\ell}\log(P_{1\ell}^0) + (1 - X_{n\ell})\log(1 - P_{1\ell}^0) \\ &+ \sum_{n \in \mathcal{N}_2 \cup \mathcal{N}_3} X_{n\ell}\log\left(\frac{P_{2\ell}^0 + P_{3\ell}^0}{2}\right) + (1 - X_{n\ell})\log\left(1 - \frac{P_{2\ell}^0 + P_{3\ell}^0}{2}\right)\Bigg],\end{aligned} \tag{22}$$

where we choose $\tilde{Q}_n^{(2)} = (1, 0) \forall n \in \mathcal{N}_1$, $\tilde{Q}_n^{(2)} = (0, 1) \forall n \in \mathcal{N}_2 \cup \mathcal{N}_3$, and $\tilde{P}_1^{(2)} = P_1^0$ and $\tilde{P}_2^{(2)} = \dfrac{P_2^0 + P_3^0}{2}$. From inequalities (21) and (22), we have

$$\begin{aligned}\widehat{\mathcal{L}}^{(3)} - \widehat{\mathcal{L}}^{(2)} \leq \frac{1}{NL}\sum_{\ell=1}^{N}\sum_{k=2,3}\sum_{n \in \mathcal{N}_k}&\left\{X_{n\ell}\left[\log(P_{k\ell}^0) - \log\left(\frac{P_{2\ell}^0 + P_{3\ell}^0}{2}\right)\right]\right. \\ &\left. + (1 - X_{n\ell})\left[\log(1 - P_{k\ell}^0) - \log\left(1 - \frac{P_{2\ell}^0 + P_{3\ell}^0}{2}\right)\right]\right\} + \delta_{NL} =: Y + \delta_{NL}.\end{aligned} \tag{23}$$

Hence,

$$\mathbb{P}\left(\widehat{\mathcal{L}}^{(3)} - \widehat{\mathcal{L}}^{(2)} \leq Y + \delta_{NL}\right) \geq 1 - (NL)^{-c}. \tag{24}$$

Now we proceed to show the concentration of $Y$ around its expected value $\mathbb{E}Y$, which is

$$\frac{1}{NL}\sum_{\ell=1}^{L}\sum_{k=2,3}\sum_{n\in\mathcal{N}_k}\left\{P^0_{k\ell}\left[\log(P^0_{k\ell}) - \log\left(\frac{P^0_{2\ell}+P^0_{3\ell}}{2}\right)\right] + (1-P^0_{k\ell})\left[\log(1-P^0_{k\ell}) - \log\left(1-\frac{P^0_{2\ell}+P^0_{3\ell}}{2}\right)\right]\right\}$$
$$= \frac{1}{3}\left(\overline{\mathrm{KL}}\left(\mathbf{P}^0_2\middle\|\frac{\mathbf{P}^0_2+\mathbf{P}^0_3}{2}\right) + \overline{\mathrm{KL}}\left(\mathbf{P}^0_3\middle\|\frac{\mathbf{P}^0_2+\mathbf{P}^0_3}{2}\right)\right)$$

since we assume $|\mathcal{N}_2| = |\mathcal{N}_3| = N/3$. By defining

$$\lambda_{k\ell} = \log\left(\frac{2P^0_{k\ell}}{P^0_{2\ell}+P^0_{3\ell}}\right) - \log\left(\frac{2-2P^0_{k\ell}}{2-(P^0_{2\ell}+P^0_{3\ell})}\right) \quad \forall k = 2,3, \ell = 1,\ldots,L,$$

we can write

$$Y - \mathbb{E}Y = \sum_{\ell=1}^{L}\sum_{k=2,3}\sum_{n\in\mathcal{N}_k}\frac{1}{NL}\lambda_{k\ell}(X_{n\ell} - P^0_{k\ell}).$$

Applying Bernstein's inequality (Lemma 9) for sequence of random variables $(Y_{n\ell})$ for $n \in \mathcal{N}_2 \cup \mathcal{N}_3$ and $\ell \in [L]$ with $Y_{n\ell} = \lambda_{k\ell}(X_{n\ell} - P^0_{k\ell})/(NL)$ having $|Y_{n\ell}| \leq 2|\lambda_{k\ell}|/(NL)$, we have

$$\mathbb{P}(Y - \mathbb{E}Y \leq t) \geq 1 - \exp\left(-\frac{t^2}{2(\sigma^2 + Mt/3)}\right), \tag{25}$$

for $M = \max_{k=2,3,\ell\in[L]} 2|\lambda_{k\ell}|/(NL)$ and

$$\sigma^2 = \frac{1}{(NL)^2}\sum_{\ell=1}^{L}\sum_{n\in\mathcal{N}_2\cup\mathcal{N}_3}\lambda^2_{k\ell}P^0_{k\ell}(1-P^0_{k\ell}) = \frac{3}{NL^2}\sum_{\ell=1}^{L}\sum_{k=2,3}\lambda^2_{k\ell}P^0_{k\ell}(1-P^0_{k\ell}).$$

Because of Assumption 2, $\log(P^0_{k\ell})$ and $\log(1-P^0_{k\ell})$ is bounded below by a constant $-\underline{c}$, so

$$\begin{aligned}\lambda_{k\ell} &= \log(P^0_{k\ell}) + \log\left(\frac{2}{P^0_{2\ell}+P^0_{3\ell}}\right) - \log(1-P^0_{k\ell}) + \log\left(1-\frac{P^0_{2\ell}+P^0_{3\ell}}{2}\right)\\ &\geq -\underline{c} + 0 + 0 - \underline{c} = -2\underline{c}.\end{aligned} \tag{26}$$

Similarly, $\lambda_{k\ell} \leq 2\underline{c}$. Therefore, $|\lambda_{k\ell}| \leq 2\underline{c}$, which implies $\sigma^2 \leq \dfrac{12\underline{c}^2}{NL}$ and $M \leq 4\dfrac{\underline{c}}{NL}$. By plugging $t = \underline{c}(\log(NL)/(NL))^{1/2}$ in (25), we can bound the exponent in the probability in the RHS as:

$$\frac{t^2}{2(\sigma^2 + Mt/3)} \geq \frac{\log(NL)}{24 + 8(\log(NL)/(NL))^{1/2}/3} \geq c_1\log(NL),$$

when $N$ and $L$ sufficiently large, where the constant $c_1$ is universal. Hence,

$$\mathbb{P}\left(Y - \mathbb{E}Y \leq \underline{c}\left(\frac{\log(NL)}{NL}\right)^{1/2}\right) \geq 1 - (NL)^{-c_1}. \tag{27}$$

Combining with (24) and using union probability bound, we have

$$\widehat{\mathcal{L}}^{(3)} - \widehat{\mathcal{L}}^{(2)} \leq \frac{1}{3}\left(\mathrm{KL}\left(\mathbf{P}_2^0 \middle\| \frac{\mathbf{P}_2^0 + \mathbf{P}_3^0}{2}\right) + \mathrm{KL}\left(\mathbf{P}_3^0 \middle\| \frac{\mathbf{P}_2^0 + \mathbf{P}_3^0}{2}\right)\right) + \delta_{NL} + \underline{c}\left(\frac{\log(NL)}{NL}\right)^{1/2}, \quad (28)$$

with probability at least $1 - (NL)^{-c} - (NL)^{-c_1}$. □

**Lemma 3.** *Assume Assumption 1 and Assumption 2. Let $\widehat{\mathcal{L}}^{(1)}$ and $\widehat{\mathcal{L}}^{(3)}$ be the maximal average log-likelihood at $K = 1$ and $K = 3$, respectively. Then, for all $N$ sufficiently large (depending on $\underline{c}$), we have*

$$\widehat{\mathcal{L}}^{(3)} - \widehat{\mathcal{L}}^{(1)} \geq \frac{1}{3}\sum_{k=1}^{3} \overline{KL}(\boldsymbol{P}_k^0 \| \tfrac{\boldsymbol{P}_1^0 + \boldsymbol{P}_2^0 + \boldsymbol{P}_3^0}{3}) - \underline{c}\left(\frac{\log(NL)}{NL}\right)^{1/2} - 2\left(\frac{\log(L)}{L}\right)^{1/2} - 2C_1\left(\frac{\log(N)}{N}\right)^{1/2}, \quad (29)$$

*with a probability exceeding $1 - (NL)^{-c_2} - 4L^{-4e^2}$, where $c_2$ and $C_1$ are universal positive constants.*

*Proof.* We first notice that the MLE for the case $K = 1$ is

$$\widehat{P}_\ell^{(1)} = \overline{X}_\ell = \sum_{n=1}^{N} X_{n\ell}/N.$$

So that

$$\begin{aligned}\widehat{\mathcal{L}}^{(1)} &= \frac{1}{NL}\sum_{n,\ell=1}^{N}\left(X_{n\ell}\log(\widehat{P}_\ell^{(1)}) + (1 - X_{n\ell})\log(1 - \widehat{P}_\ell^{(1)})\right) \\ &= \frac{1}{L}\sum_{\ell=1}^{L}\left(\overline{X}_\ell \log(\overline{X}_\ell) + (1 - \overline{X}_\ell)\log(1 - \overline{X}_\ell)\right).\end{aligned}$$

Let $\overline{P}_\ell = \frac{1}{3}(P_{1\ell}^0 + P_{2\ell}^0 + P_{3\ell}^0)$ for all $\ell \in [L]$. By Lemma 5, we have

$$\left|\frac{1}{L}\sum_{\ell}\left(\overline{X}_\ell \log(\overline{X}_\ell) - \overline{P}_\ell \log(\overline{P}_\ell)\right)\right| \leq \left(\frac{\log(L)}{L}\right)^{1/2} + C_1\left(\frac{\log(N)}{N}\right)^{1/2}, \quad (30)$$

with probability exceeding $1 - 2L^{-4e^2}$, for some universal constant $C_1$. Similarly, we have

$$\left|\frac{1}{L}\sum_{\ell}\left((1 - \overline{X}_\ell)\log(1 - \overline{X}_\ell) - (1 - \overline{P}_\ell)\log(1 - \overline{P}_\ell)\right)\right| \leq \left(\frac{\log(L)}{L}\right)^{1/2} + C_1\left(\frac{\log(N)}{N}\right)^{1/2}, \quad (31)$$

with probability exceeding $1 - 2L^{-4e^2}$. Combining them and using Bonferroni's Inequality, we have

$$\left|\widehat{\mathcal{L}}^{(1)} - \frac{1}{L}\sum_{\ell}\left(\overline{P}_\ell \log(\overline{P}_\ell) + (1 - \overline{P}_\ell)\log(1 - \overline{P}_\ell)\right)\right| \leq 2\left(\frac{\log(L)}{L}\right)^{1/2} + 2C_1\left(\frac{\log(N)}{N}\right)^{1/2}, \quad (32)$$

with probability exceeding $1 - 4L^{-4e^2}$. Moreover, by Lemma 4, we have

$$\widehat{\mathcal{L}}^{(3)} \geq \frac{1}{3}\sum_{k=1}^{3}\frac{1}{L}\sum_{\ell=1}^{L}\left(P_{k\ell}^0 \log(P_{k\ell}^0) + (1 - P_{k\ell}^0)\log(1 - P_{k\ell}^0)\right) - \underline{c}\left(\frac{\log(NL)}{NL}\right)^{1/2}, \quad (33)$$

with probability exceeding $1-(NL)^{-c_2}$. Hence, in the union of those two events, we have

$$\begin{aligned}\widehat{\mathcal{L}}^{(3)}-\widehat{\mathcal{L}}^{(1)} &\geq \frac{1}{3}\sum_{k=1}^{3}\frac{1}{L}\sum_{\ell=1}^{L}\left(P_{k\ell}^{0}(\log(P_{k\ell}^{0})-\log(\overline{P}_{\ell}))+(1-P_{k\ell}^{0})(\log(1-P_{k\ell}^{0})-\log(1-\overline{P}_{\ell}))\right)\\ &\quad -\underline{c}\left(\frac{\log(NL)}{NL}\right)^{1/2}-2\left(\frac{\log(L)}{L}\right)^{1/2}-2C_1\left(\frac{\log(N)}{N}\right)^{1/2}\\ &=\frac{1}{3}\sum_{k=1}^{3}\overline{\mathrm{KL}}(\mathbf{P}_k^0\|\tfrac{\mathbf{P}_1^0+\mathbf{P}_2^0+\mathbf{P}_3^0}{3})-\underline{c}\left(\frac{\log(NL)}{NL}\right)^{1/2}-2\left(\frac{\log(L)}{L}\right)^{1/2}-2C_1\left(\frac{\log(N)}{N}\right)^{1/2},\end{aligned}$$

with a probability exceeding $1-(NL)^{-c_2}-4L^{-4e^2}$. □

Lemma 3 uses results from Lemma 4 and Lemma 5, which we now state and prove.

**Lemma 4.** *We have*

$$\widehat{\mathcal{L}}^{(3)} \geq \frac{1}{3}\sum_{k=1}^{3}\frac{1}{L}\sum_{\ell=1}^{L}\left(P_{k\ell}^{0}\log(P_{k\ell}^{0})+(1-P_{k\ell}^{0})\log(1-P_{k\ell}^{0})\right)-\underline{c}\left(\frac{\log(NL)}{NL}\right)^{1/2}, \tag{34}$$

*with probability exceeding* $1-(NL)^{-c_2}$ *for a universal constant* $c_2>0$.

*Proof.* Firstly, because $\widehat{\mathcal{L}}^{(3)}$ is the maximal average log-likelihood when $K=3$, we have that it is no smaller than the average log-likelihood of the model with true parameters, i.e.,

$$\widehat{\mathcal{L}}^{(3)} \geq \mathcal{L}(\mathbf{Q}^0,\mathbf{P}^0)=\frac{1}{NL}\sum_{k=1}^{3}\sum_{n\in\mathcal{N}_k}\sum_{\ell=1}^{L}\left(X_{n\ell}\log(P_{k\ell}^{0})+(1-X_{n\ell})\log(1-P_{k\ell}^{0})\right)=:Y'. \tag{35}$$

Because $|\log(P_{k\ell}^0)|\leq\underline{c}$ and $\log(1-P_{k\ell}^0)\leq\underline{c}$, we can use the same technique as bounding the concentration of $Y$ around its mean as in Lemma 2 and arrive at the concentration bound for $Y'$:

$$\mathbb{P}\left(Y'-\mathbb{E}Y'\geq-\underline{c}\left(\frac{\log(NL)}{NL}\right)^{1/2}\right)\geq 1-(NL)^{-c_2},$$

for a universal constant $c_2$. Combining the two inequalities above and noticing that

$$\mathbb{E}Y'=\frac{1}{3}\sum_{k=1}^{3}\frac{1}{L}\sum_{\ell=1}^{L}\left(P_{k\ell}^{0}\log(P_{k\ell}^{0})+(1-P_{k\ell}^{0})\log(1-P_{k\ell}^{0})\right),$$

yields the conclusion of this lemma. □

**Lemma 5.** *Let* $\overline{X}_\ell=\sum_{n=1}^{N}X_{n\ell}/N$ *be the estimate of the allele frequency for the admixture model with* $K=1$, *then for all sufficiently large* $N$ *(depending on* $\underline{c}$*), we have with probability exceeding* $1-2L^{-4e^2}$ *that*

$$\left|\frac{1}{L}\sum_{\ell}\left(\overline{X}_\ell\log(\overline{X}_\ell)-\overline{P}_\ell\log(\overline{P}_\ell)\right)\right|\leq\left(\frac{\log(L)}{L}\right)^{1/2}+C_1\left(\frac{\log(N)}{N}\right)^{1/2}, \tag{36}$$

*where* $\overline{P}_\ell=\frac{1}{3}(P_{1\ell}^0+P_{2\ell}^0+P_{3\ell}^0)$, *and* $C_1$ *is a universal constant.*

*Proof of Lemma 5.* An application of the triangle inequality yields:

$$\frac{1}{L}\left|\sum_{\ell=1}^{L}\left(\overline{X}_\ell\log(\overline{X}_\ell)-\overline{P}_\ell\log(\overline{P}_\ell)\right)\right| \le \frac{1}{L}\left|\sum_{\ell=1}^{L}\left(\overline{X}_\ell\log(\overline{X}_\ell)-\mathbb{E}\overline{X}_\ell\log(\overline{X}_\ell)\right)\right| + \left(\frac{1}{L}\sum_{\ell=1}^{L}\left|\mathbb{E}\overline{X}_\ell\log(\overline{X}_\ell)-\overline{P}_\ell\log(\overline{P}_\ell)\right|\right). \tag{37}$$

Hence, it suffices to bound the two terms in the RHS and then derive the desired concentration. We divide them into two steps.

**Step 1. Concentration of $\sum \overline{X}_\ell\log(\overline{X}_\ell)$ around its mean.** The function $x\mapsto x\log(x)$ is bounded in $[-1/e,0]$ when $x\in[0,1]$. Hence, the random variable

$$\overline{X}_\ell\log(\overline{X}_\ell)-\mathbb{E}\overline{X}_\ell\log(\overline{X}_\ell)$$

is mean zero, bounded (in $[-1/e,1/e]$) for all $\ell\in[L]$. An application of Hoeffding's inequality gives:

$$\mathbb{P}\left(\left|\frac{1}{L}\sum_{\ell}\left(\overline{X}_\ell\log(\overline{X}_\ell)-\mathbb{E}\overline{X}_\ell\log(\overline{X}_\ell)\right)\right|\ge t\right)\le 2e^{-4Le^2t^2}, \tag{38}$$

for all $t\ge 0$. Plugging in $t=(\log L/L)^{1/2}$, we have

$$\mathbb{P}\left(\left|\frac{1}{L}\sum_{\ell}\left(\overline{X}_\ell\log(\overline{X}_\ell)-\mathbb{E}\overline{X}_\ell\log(\overline{X}_\ell)\right)\right|\ge\left(\frac{\log L}{L}\right)^{1/2}\right)\le 2L^{-4e^2}, \tag{39}$$

**Step 2. Bound $|\mathbb{E}\overline{X}_\ell\log(\overline{X}_\ell)-\overline{P}_\ell\log(\overline{P}_\ell)|$.** Apply Chernoff's inequality for small deviation of Bernoulli random variables (Lemma 8), we have

$$\mathbb{P}(|\overline{X}_\ell-\overline{P}_\ell|\ge\delta\overline{P}_\ell)\le 2e^{-N\overline{P}_\ell\delta^2/3}, \tag{40}$$

for all $\delta\in(0,1]$ and $\ell\in[L]$. In the event $\{|\overline{X}_\ell-\overline{P}_\ell|\le\delta\overline{P}_\ell)\}$, we have $\overline{X}_\ell\in[\overline{P}_\ell(1-\delta),\overline{P}_\ell(1+\delta)]$, so that

$$\log(\overline{P}_\ell)+\log(1-\delta)\le\log(\overline{X}_\ell)\le\log(\overline{P}_\ell)+\log(1+\delta),$$

which implies

$$\begin{aligned}|\overline{X}_\ell\log(\overline{X}_\ell)-\overline{P}_\ell\log(\overline{P}_\ell)| &\le |\overline{X}_\ell-\overline{P}_\ell||\log(\overline{P}_\ell)|+\overline{X}_\ell|\log(\overline{X}_\ell)-\log(\overline{P}_\ell)|\\ &\le \delta\overline{P}_\ell|\log(\overline{P}_\ell)|+(1+\delta)\overline{P}_\ell\max\{-\log(1-\delta),\log(1+\delta)\}\\ &\le \delta\overline{P}_\ell|\log(\overline{P}_\ell)|+(-\log(1-\delta))(1+\delta)\overline{P}_\ell,\end{aligned} \tag{41}$$

because $\log(1+\delta)\le-\log(1-\delta)$ for all $\delta\in(0,1]$. Hence,

$$\begin{aligned}|\mathbb{E}\overline{X}_\ell\log(\overline{X}_\ell)-\overline{P}_\ell\log(\overline{P}_\ell)| &\le \mathbb{E}\left[|\overline{X}_\ell\log(\overline{X}_\ell)-\overline{P}_\ell\log(\overline{P}_\ell)|1_{\{|\overline{X}_\ell-\overline{P}_\ell|\le\delta\overline{P}_\ell)\}}\right]\\ &\quad+\mathbb{E}\left[|\overline{X}_\ell\log(\overline{X}_\ell)-\overline{P}_\ell\log(\overline{P}_\ell)|1_{\{|\overline{X}_\ell-\overline{P}_\ell|\ge\delta\overline{P}_\ell)\}}\right]\\ &\le \delta\overline{P}_\ell|\log(\overline{P}_\ell)|+(-\log(1-\delta))(1+\delta)\overline{P}_\ell\end{aligned}$$

$$+ (1/e) \times \mathbb{P}(|\overline{X}_\ell - \overline{P}_\ell| \geq \delta \overline{P}_\ell)$$
$$\leq \delta \overline{P}_\ell |\log(\overline{P}_\ell)| + (-\log(1-\delta))(1+\delta)\overline{P}_\ell + 2e^{-N\overline{P}_\ell \delta^2/3 - 1}.$$

Choose $\delta = \left(\frac{3\log(N)}{\overline{P}_\ell N}\right)^{1/2} \leq \left(\frac{3\log(N)e^{\underline{c}}}{N}\right)^{1/2} \to 0$ as $N \to \infty$, we have

$$\delta \overline{P}_\ell |\log(\overline{P}_\ell)| = \left(\frac{3\log(N)}{N}\right)^{1/2} \overline{P}_\ell^{1/2} |\log(\overline{P}_\ell)| \leq \frac{2}{e}\left(\frac{3\log(N)}{N}\right)^{1/2},$$

since $x^{1/2}\log(x) \in [-2/e, 0]$ for all $x \in [0,1]$. Moreover, when $N$ is sufficiently large so that $\delta < 0.8$, we have $-\log(1-\delta) \leq 2\delta$, which implies

$$(-\log(1-\delta))(1+\delta)\overline{P}_\ell \leq 4\delta \overline{P}_\ell \leq 4\left(\frac{3\log(N)}{N}\right)^{1/2}.$$

Finally,

$$2e^{-N\overline{P}_\ell \delta^2/3 - 1} = \frac{2}{e}\frac{1}{\sqrt{N}}.$$

Hence,

$$|\mathbb{E}\overline{X}_\ell \log(\overline{X}_\ell) - \overline{P}_\ell \log(\overline{P}_\ell)| \leq C_1 \left(\frac{\log(N)}{N}\right)^{1/2},$$

for a universal constant $C_1$ for all sufficiently large $N$. □

##### 1.3 Empirical Process theory and the proof of Lemma 1

Lemma 1 is proved using the Empirical Process framework applied for M-estimation [van de Geer, 2000], of which some main notations and results are recalled here. In this subsection, we fix an over-fitted level $K \geq K_0$ and will setup and prove Lemma 1 for this particular $K$.

Let $\mathcal{X} = \{0,1\}$ be the support (possible value) of genotype data. Given two sequences of non-negative functions $(F_{n\ell})_{n,\ell=1}^{N,L}$ and $(F'_{n\ell})_{n,\ell=1}^{N,L}$ on $\mathcal{X}^{N\times L}$, define the Hellinger process distance [van de Geer, 2000] between product functions $F = \otimes_{n,\ell} F_{n\ell}$ and $F' = \otimes_{n,\ell} F'_{n\ell}$ (and between each pair of elements) as

$$H^2(F, F') := \sum_{n,\ell=1}^{N,L} H^2(F_{n\ell}, F'_{n\ell}), \qquad H^2(F_{n\ell}, F'_{n\ell}) := \frac{1}{2}\sum_{x=0}^{1}\left(\sqrt{F_{n\ell}(x)} - \sqrt{F'_{n\ell}(x)}\right)^2. \tag{42}$$

and the average Hellinger process is defined as

$$\overline{H}^2(F, F') := \frac{1}{NL} H^2(F, F'). \tag{43}$$

Let $\mathcal{A} = \{(\mathbf{Q}, \mathbf{P}) : \mathbf{Q} = (Q_{nk})_{n,k} \subset [0,1], \sum_{k=1}^K Q_{nk} = 1, \mathbf{P} = (P_{k\ell})_{k,\ell} \subset [0,1]\}$ be the the parameter space of the admixture model for $N$ haploids at $L$ loci and $K$ populations. For a set of parameters $(\mathbf{Q}, \mathbf{P}) \in \mathcal{A}$, let $F^{(\mathbf{Q},\mathbf{P})} = (F_{n\ell}^{(\mathbf{Q},\mathbf{P})})_{n,\ell=1}^{N,L}$ be the collection of p.m.f. of $X_{n\ell}$ in the admixture model with corresponding parameters $\mathbf{P}$ and $\mathbf{Q}$, i.e.,

$$F_{n\ell}^{(\mathbf{Q},\mathbf{P})}(x) = \mathrm{Ber}(x|\sum_{k=1}^K Q_{nk}P_{k\ell}), \quad \forall n \in [N], \ell \in [L].$$

Given true parameters $\mathbf{P}^0 \in [0,1]^{K_0\times L}$ and $\mathbf{Q}^0 \in [0,1]^{N\times K_0}$, let $\Theta^0 = (\theta^0_{n\ell})_{n,\ell=1}^{N,L} = \mathbf{Q}^0\mathbf{P}^0$, i.e, of which elements are calculated by $\theta^0_{n\ell} = \sum_{k=1}^{K_0} Q^0_{nk}P^0_{k\ell}, \quad n \in [N], \ell \in [L]$. The true probability function $F^0 = \otimes_{n,\ell}F^0_{n\ell}$ is defined by

$$F^0_{n\ell}(x) := \mathrm{Ber}(x|\theta^0_{n\ell}) = \begin{cases} \theta^0_{n\ell}, & x = 1, \\ 1-\theta^0_{n\ell}, & x = 0. \end{cases} \tag{44}$$

Let the estimated individual's allele frequency be $\widehat{\theta}_{n\ell} = \sum_{k=1}^{K} \widehat{Q}_{nk}\widehat{P}_{k\ell}$. This results in the estimated probability function $\widehat{F} = \otimes\widehat{F}_{n\ell}$ defined by

$$\widehat{F}_{n\ell}(x) := \mathrm{Ber}(x|\widehat{\theta}_{n\ell}) = \begin{cases} \widehat{\theta}_{n\ell}, & x = 1, \\ 1-\widehat{\theta}_{n\ell}, & x = 0. \end{cases} \tag{45}$$

**Covering number and entropy number.** The Empirical process theory provides many useful concentration inequalities in the space of probability functions $(F_{n\ell})$ so that the convergence rate of MLE boils down to calculate (or sufficiently provide an upper bound for) the smallest number of balls to cover this space in the Hellinger process distance. This number is often referred to as the "covering number", which we will now define.

**Definition 1.** *For $\delta > 0$ and a space $\mathcal{F}$ of product p.m.f.'s $F = \otimes_{n,\ell}F_{n\ell}$ on $\mathcal{X}^{N\times L}$, let $\mathcal{N}_B(\delta, \mathcal{F})$ be the smallest integer $M$ such that there exists a collection of non-negative functions $\{\underline{\mathbf{F}}^{(j)}, \overline{\mathbf{F}}^{(j)}\}_{j=1}^M$ with $\underline{\mathbf{F}}^{(j)} = (\underline{F}^{(j)}_{n\ell})_{n,\ell=1}^{N,L}$ and $\overline{\mathbf{F}}^{(j)} = (\overline{F}^{(j)}_{n\ell})_{n,\ell=1}^{N,L}$ such that for every $F \in \mathcal{F}$, there is an index $j \in [M]$ such that*

*(i)* $\overline{H}\left(\dfrac{\underline{\mathbf{F}}^{(j)} + \overline{\mathbf{F}}^0}{2}, \dfrac{\overline{\mathbf{F}}^{(j)} + \mathbf{F}^0}{2}\right) \le \delta$ *and*

*(ii)* $\underline{F}^{(j)}_{n\ell}(x) \le F_{n\ell}(x) \le \overline{F}^{(j)}_{n\ell}(x)$ *for all $x \in \mathcal{X}$ and $n \in [N], \ell \in [L]$.*

*Then $\mathcal{N}_B(\delta, \mathcal{F})$ and $\mathcal{H}_B(\delta, \mathcal{F}) = \log \mathcal{N}_B(\delta, \mathcal{F})$ are called the Hellinger covering number and entropy number with bracketing, respectively.*

For $c_0$ being a suitable universal constant, define the entropy integral:

$$\mathcal{J}_B(\delta) := \int_{\delta^2/c_0}^{\delta} \mathcal{H}_B^{1/2}(u, \mathcal{F})du \vee \delta, \quad 0 < \delta \le 1, \tag{46}$$

where $a \vee b$ denotes the maximum of two numbers $a$ and $b$.

The two main results, of which notation is adapted to our model, are stated as follows.

**Theorem A** (Theorem 8.14 in [van de Geer, 2000]). *Take function*

$$\Psi(\delta) \ge \mathcal{J}_B\left(\delta, \left\{F^{(\boldsymbol{Q},\boldsymbol{P})} : (\boldsymbol{Q}, \boldsymbol{P}) \in \mathcal{A}\right\}\right)$$

*in such a way that $\Psi(\delta)/\delta^2$ is a non-increasing function of $\delta$. Then for a constant $c$ satisfying*

$$\sqrt{NL}\delta^2_{NL} \ge c\Psi(\delta_{NL}), \tag{47}$$

*we have that for all $\delta \ge \delta_{NL}$,*

$$\mathbb{P}\left(\overline{H}\left(\widehat{F}, F^0\right) \ge \delta\right) \le c\exp\left(-\frac{NL\delta^2}{c^2}\right). \tag{48}$$

**Theorem B** (Theorem 8.13 in [van de Geer, 2000]). *Let positive numbers $R, D, C_1, a$, and a subset of parameter space $\mathcal{B} \subset \mathcal{A}$ satisfy*

$$\overline{H}(F^{(\boldsymbol{Q},\boldsymbol{P})}, F^0) \leq R \quad \forall(\boldsymbol{Q},\boldsymbol{P}) \in \mathcal{B}, \tag{49}$$

$$b \leq C_1 \min\{\sqrt{NL}R^2, 8\sqrt{NL}R\}, \tag{50}$$

*and*

$$b \geq \sqrt{D^2(C_1+1)} \left( \int_{b/(2^6\sqrt{NL})}^{R} \mathcal{H}_B^{1/2} \left( \frac{u}{\sqrt{2}}, \left\{ F^{(\boldsymbol{Q},\boldsymbol{P})} : (\boldsymbol{Q},\boldsymbol{P}) \in \mathcal{B} \right\} \right) du \vee R \right), \tag{51}$$

*then*

$$\mathbb{P}\Big( \sup_{(\boldsymbol{Q},\boldsymbol{P})\in\mathcal{B}} \sqrt{NL} |\bar{Z}^{(\boldsymbol{Q},\boldsymbol{P})} - \bar{A}^{(\boldsymbol{Q},\boldsymbol{P})}| \geq b \Big) \leq D \exp\left[ -\frac{b^2}{D^2(C_1+1)R^2} \right], \tag{52}$$

*where*

$$Z_{n\ell}^{(\boldsymbol{Q},\boldsymbol{P})} = \frac{1}{2} \log\left( \frac{\mathrm{Ber}(x_{n\ell}|\theta_{n\ell}) + \mathrm{Ber}(x_{n\ell}|\theta^0_{n\ell})}{2\mathrm{Ber}(x_{n\ell}|\theta^0_{n\ell})} \right), \quad \theta_{n\ell} = \sum_k Q_{nk}P_{k\ell},$$

*and*

$$\bar{Z}^{(\boldsymbol{Q},\boldsymbol{P})} = \frac{1}{NL} \sum_{n,\ell}^{N,L} Z_{n\ell}^{(\boldsymbol{Q},\boldsymbol{P})}, \quad \bar{A}^{(\boldsymbol{Q},\boldsymbol{P})} = \frac{1}{NL} \sum_{n,\ell=1}^{N,L} \mathbb{E}_{X_{n\ell}\sim\mathrm{Ber}(\theta^0_{n\ell})} Z_{n\ell}^{(\boldsymbol{Q},\boldsymbol{P})}.$$

*Proof of Lemma 1.* Similar to the proof of Theorem 6.3.2 in [Do, 2025] but applying to haploid data (Bernoulli model) instead of diploid data (Binomial model), we can bound the entropy number of the admixture models' space as

$$\mathcal{H}_B\left(\delta, \left\{ F^{(\mathbf{Q},\mathbf{P})} : (\mathbf{Q},\mathbf{P}) \in \mathcal{A} \right\}\right) \leq C_0 K(N+L)\log(1/\delta), \tag{53}$$

for a universal constant $C_0$ and all sufficiently small $\delta$, and with the choice

$$\delta_{NL} = C \left( \frac{K(N+L)\log(NL)}{NL} \right)^{1/2},$$

the convergence of MLE (as implied from Theorem A) is

$$\mathbb{P}\left( \overline{H}\left(\widehat{F}, F^0\right) \leq C \left( \frac{K(N+L)\log(NL)}{NL} \right)^{1/2} \right) \geq 1 - c_1(NL)^{-1/c_1^2}. \tag{54}$$

Now apply Theorem B for $\mathcal{B} = \{(\mathbf{Q},\mathbf{P}) : \overline{H}(F^{(\mathbf{Q},\mathbf{P})}, F^0) \leq R\}$, $R = \delta_{NL}$, $b = \sqrt{NL}\delta^2_{NL} = C^2K(N+L)\log(NL)/\sqrt{NL}$ and $C_1 = 1$, we readily obtain that (50) is correct for all sufficiently large $N$ and $L$. Inequality (51) also holds since $b \geq R$ and

$$b \gtrsim R \log\left( \frac{2^6\sqrt{NL}}{b} \right) \gtrsim \int_{b/(2^6\sqrt{NL})}^{R} \mathcal{H}_B^{1/2}\left( \frac{u}{\sqrt{2}}, \left\{ F^{(\mathbf{Q},\mathbf{P})} : (\mathbf{Q},\mathbf{P}) \in \mathcal{B} \right\} \right) du,$$

which we apply (53) and the fact that $\log(1/\delta)$ is a decreasing function. Hence,

$$\mathbb{P}\Big(\sup_{(\mathbf{Q},\mathbf{P})\in\mathcal{B}}|\bar{Z}^{(\mathbf{Q},\mathbf{P})}-\bar{A}^{(\mathbf{Q},\mathbf{P})}|\leq C^2\frac{K(N+L)\log(NL)}{NL}\Big)\geq 1-D(NL)^{-D_1}, \tag{55}$$

for some constant $D_1$. Combining the two events in (54) and (55) we have with the probability at least $1-c_1(NL)^{-1/c_1^2}-D(NL)^{-D_1}$ that

$$\overline{Z}^{(\widehat{\mathbf{Q}},\widehat{\mathbf{P}})}-\overline{A}^{(\widehat{\mathbf{Q}},\widehat{\mathbf{P}})}\leq C^2\frac{K(N+L)\log(NL)}{NL}. \tag{56}$$

Moreover,

$$\overline{A}^{(\widehat{\mathbf{Q}},\widehat{\mathbf{P}})}=-\frac{1}{NL}\sum_{n,\ell}\mathrm{KL}\left(\mathrm{Ber}(\theta^0_{n\ell}),\mathrm{Ber}\left(\tfrac{\theta^0_{n\ell}+\widehat{\theta}_{n\ell}}{2}\right)\right)\leq 0,$$

and by the concavity of the log function,

$$\bar{Z}^{(\widehat{\mathbf{Q}},\widehat{\mathbf{P}})}=\frac{1}{2NL}\sum_{n,\ell}^{N,L}Z_{n\ell}^{(\widehat{\mathbf{Q}},\widehat{\mathbf{P}})}\geq\frac{1}{NL}\sum_{n,\ell}^{N,L}\log\left(\frac{\mathrm{Ber}(x_{n\ell}|\widehat{\theta}_{n\ell})}{\mathrm{Ber}(x_{n\ell}|\theta^0_{n\ell})}\right)=\widehat{\mathcal{L}}^{(K)}-\mathcal{L}(\mathbf{Q}^0,\mathbf{P}^0).$$

Hence,

$$\widehat{\mathcal{L}}^{(K)}\leq\mathcal{L}(\mathbf{Q}^0,\mathbf{P}^0)+\frac{C^2K(N+L)\log(NL)}{NL}$$

with probability at least $1-(NL)^{-c}$ with a universal constant $c$ (derived from $c_1,D,D_1$). Writing $C^2$ as $C$ instead, we have the upper bound part of Lemma 1. The lower bound is trivial from the definition of the MLE and the fact that $K\geq K_0$. □

### 2 Theorem 2

Here is the formal version of Theorem 2 in the main text:

**Theorem 2** (Small-drift expansion of $\mathcal{D}_{31}$ and $\mathcal{D}_{32}$ under nested BN)**.** *Consider the nested Balding–Nichols model (root frequency $P^*_\ell$, internal node $P^0_{23\ell}$, and leaves $P^0_{1\ell},P^0_{2\ell},P^0_{3\ell}$) with parameters $F_{root},F_{sub},F_{out}$, and assume $\varepsilon\leq P^*_\ell\leq 1-\varepsilon$ for all $\ell$ for some fixed $\varepsilon\in(0,1/2)$. let*

$$m_\ell:=\frac{P^0_{1\ell}+P^0_{2\ell}+P^0_{3\ell}}{3},\qquad m_{23,\ell}:=\frac{P^0_{2\ell}+P^0_{3\ell}}{2}.$$

*Then, as $(F_{root},F_{sub},F_{out})\to 0$,*

$$\mathbb{E}[\mathcal{D}_{32}]=\frac{1}{2}F_{sub}(1-F_{root})+o(F_{root}+F_{sub}+F_{out}), \tag{57}$$

$$\mathbb{E}\left[\frac{1}{3}\mathcal{D}_{31}\right]=\frac{1}{9}(F_{out}+F_{root})+\frac{2}{9}F_{sub}(1-F_{root})+o(F_{root}+F_{sub}+F_{out}). \tag{58}$$

*In particular, under the variance-matching choice $F_{out}=F_{sub}+F_{root}-F_{sub}F_{root}$,*

$$\mathbb{E}\left[\frac{1}{3}\mathcal{D}_{31}-\mathcal{D}_{32}\right]=\frac{2}{9}F_{root}-\frac{1}{6}F_{sub}(1-F_{root})+o(F_{root}+F_{sub}), \tag{59}$$

*so in the diffusion regime where $F_{root},F_{sub}\to 0$ the sign is governed by $4F_{root}-3F_{sub}$ (equivalently $F_{root}/F_{sub}\gtrless 3/4$).*

#### 2.1 Supporting lemmas

**Lemma 6** (Local quadratic approximation of Bernoulli KL)**.** *Fix* $\varepsilon \in (0, 1/2)$*. There exists a constant* $C_\varepsilon > 0$ *such that for all* $p, q \in [\varepsilon, 1-\varepsilon]$*,*

$$KL(\mathrm{Ber}(p)\|\mathrm{Ber}(q)) = \frac{(p-q)^2}{2q(1-q)} + R(p,q), \tag{60}$$

*where the remainder satisfies* $|R(p,q)| \le C_\varepsilon |p-q|^3$*.*

*Proof.* Let $\phi(x) = x\log x + (1-x)\log(1-x)$ on $(0,1)$. Then $\mathrm{KL}(\mathrm{Ber}(p)\|\mathrm{Ber}(q)) = \phi(p) - \phi(q) - \phi'(q)(p-q)$. By Taylor's theorem, there exists $\xi$ between $p$ and $q$ such that

$$\mathrm{KL}(\mathrm{Ber}(p)\|\mathrm{Ber}(q)) = \frac{1}{2}\phi''(q)(p-q)^2 + \frac{1}{6}\phi'''(\xi)(p-q)^3.$$

Since $\phi''(x) = 1/(x(1-x))$ and $\phi'''(x) = (2x-1)/(x^2(1-x)^2)$, we have $\sup_{x\in[\varepsilon,1-\varepsilon]} |\phi'''(x)| < \infty$, yielding (60). □

We need an additional lemma to bound the probability of the sub-population's allele frequency away from 0 under the Balding–Nichols model.

**Lemma 7** (Boundary avoidance under Balding–Nichols)**.** *Fix* $0 < \varepsilon_0 < \varepsilon < 1/2$*. There exist constants* $c, C > 0$ *(depending only on* $\varepsilon, \varepsilon_0$*) such that the following holds for all sufficiently small* $F \in (0,1)$*: for every* $x \in [\varepsilon, 1-\varepsilon]$*, if*

$$Y \mid x \sim \mathrm{Beta}\left(\frac{1-F}{F}x, \frac{1-F}{F}(1-x)\right),$$

*then*

$$\mathbb{P}(Y \notin [\varepsilon_0, 1-\varepsilon_0] \mid x) \le C\exp\left(-\frac{c}{F}\right).$$

*Proof.* We bound the lower tail; the upper tail follows by the same argument applied to $1-Y$. Write $\alpha = (1-F)/F$ and note $\alpha \to \infty$ as $F \to 0$. Conditional on $x$, the density of $Y$ is

$$f_{x,\alpha}(y) = \frac{y^{\alpha x-1}(1-y)^{\alpha(1-x)-1}}{B(\alpha x, \alpha(1-x))} =: \frac{g(y)}{B(\alpha x, \alpha(1-x))} \quad y \in (0,1).$$

Then

$$\frac{d}{dy}\log g(y) = \frac{\alpha x - 1}{y} - \frac{\alpha(1-x)-1}{1-y} = \frac{\alpha(x-y) - 1 + 2y}{y(1-y)}.$$

For $y \in (0, \varepsilon_0]$ we have $x - y \ge \varepsilon - \varepsilon_0 > 0$, so for all sufficiently small $F$ (equivalently, all sufficiently large $\alpha$) the derivative is strictly positive on $(0, \varepsilon_0]$. Hence $g$ is increasing on $(0, \varepsilon_0]$, and therefore for all $y \in (0, \varepsilon_0]$ we have $g(y) \le g(\varepsilon_0)$. Thus,

$$\mathbb{P}(Y \le \varepsilon_0 \mid x) \le \frac{\varepsilon_0 g(\varepsilon_0)}{B(\alpha x, \alpha(1-x))} = \frac{\varepsilon_0^{\alpha x}(1-\varepsilon_0)^{\alpha(1-x)-1}}{B(\alpha x, \alpha(1-x))}. \tag{61}$$

Note that $B(\alpha x, \alpha(1-x)) = \Gamma(\alpha x)\Gamma(\alpha(1-x))/\Gamma(\alpha)$, recall the following inequalities related to Stirling's approximation (a detailed derivation can be seen in [Robbins, 1955]):

$$\sqrt{2\pi z}\left(\frac{z}{e}\right)^z < \Gamma(z+1) < 1.1 * \sqrt{2\pi z}\left(\frac{z}{e}\right)^z \quad \forall z \ge 1.$$

Applying this result, we have

$$\Gamma(\alpha) = \frac{\Gamma(\alpha+1)}{\alpha} < 1.1 * \frac{1}{\alpha}\sqrt{2\pi\alpha}\left(\frac{\alpha}{e}\right)^{\alpha} = 1.1 * \sqrt{\frac{2\pi}{\alpha}}\left(\frac{\alpha}{e}\right)^{\alpha},$$

and

$$\Gamma(\alpha x) = \frac{\Gamma(\alpha x+1)}{\alpha x} > \sqrt{\frac{2\pi}{\alpha x}}\left(\frac{\alpha x}{e}\right)^{\alpha x},\ \Gamma(\alpha(1-x)) > \sqrt{\frac{2\pi}{\alpha(1-x)}}\left(\frac{\alpha(1-x)}{e}\right)^{\alpha(1-x)}.$$

Note that those inequalities are valid when $\alpha x$ and $\alpha(1-x) \geq 1$, which is equivalent to $F \leq \frac{x}{x+1} \leq \epsilon$ (i.e., for all sufficiently small $F$). Combining them yields

$$\frac{1}{B(\alpha x, \alpha(1-x))} \leq C\sqrt{\alpha}\,\frac{1}{x^{\alpha x}(1-x)^{\alpha(1-x)}}$$

for all $\alpha$ large enough and all $x \in [\varepsilon, 1-\varepsilon]$. Together with inequality (61), we have

$$\mathbb{P}(Y \leq \varepsilon_0 \mid x) \leq C\sqrt{\alpha}\exp\Big(-\alpha\,\mathrm{KL}(\mathrm{Ber}(x)\|\mathrm{Ber}(\varepsilon_0))\Big),$$

since $\mathrm{KL}(\mathrm{Ber}(x)\|\mathrm{Ber}(\varepsilon_0)) = x\log\big(x/\varepsilon_0\big) + (1-x)\log\big((1-x)/(1-\varepsilon_0)\big)$. Because $x \mapsto \mathrm{KL}(\mathrm{Ber}(x)\|\mathrm{Ber}(\varepsilon_0))$ is continuous and strictly positive on $[\varepsilon, 1-\varepsilon]$ (as $\varepsilon_0 < \varepsilon < x$), its infimum over this interval is a positive constant $\kappa > 0$. Hence

$$\mathbb{P}(Y \leq \varepsilon_0 \mid x) \leq C\sqrt{\alpha}\,e^{-\kappa\alpha}.$$

Finally, since $\log\alpha = o(\alpha)$ as $\alpha \to \infty$, we have $\sqrt{\alpha} \leq e^{(\kappa/2)\alpha}$ for all sufficiently large $\alpha$, and thus

$$C\sqrt{\alpha}\,e^{-\kappa\alpha} \leq Ce^{-(\kappa/2)\alpha}.$$

Recalling $\alpha = (1-F)/F$, for $F \leq 1/2$ we have $\alpha \geq 1/(2F)$, so $e^{-(\kappa/2)\alpha} \leq \exp(-\kappa/(4F))$. Therefore

$$\mathbb{P}(Y \leq \varepsilon_0 \mid x) \leq Ce^{-c/F}$$

for some $c > 0$ and all sufficiently small $F$. □

#### 2.2 Proof of the theorem

We work locus-by-locus and suppress the index $\ell$. Fix any $\varepsilon_0 \in (0, \varepsilon)$ and $x \in [\varepsilon, 1-\varepsilon]$, working conditional on $P^* = x$. Define the *good event* at this locus

$$G := E_{\varepsilon_0} = \{\varepsilon_0 \leq P_k^0 \leq 1-\varepsilon_0\ \forall k \in [3]\}, \qquad B := G^c.$$

Since $m_{23} = (P_2^0 + P_3^0)/2$ and $m = (P_1^0 + P_2^0 + P_3^0)/3$, on $G$ we have $\varepsilon_0 \leq m_{23}, m \leq 1-\varepsilon_0$ as well.

We first split expectations into good/bad parts:

$$\mathbb{E}[\mathcal{D}_{32} \mid x] = \mathbb{E}[\mathcal{D}_{32}\mathbf{1}_G \mid x] + \mathbb{E}[\mathcal{D}_{32}\mathbf{1}_B \mid x], \qquad \mathbb{E}[\mathcal{D}_{31} \mid x] = \mathbb{E}[\mathcal{D}_{31}\mathbf{1}_G \mid x] + \mathbb{E}[\mathcal{D}_{31}\mathbf{1}_B \mid x].$$

On $B$ we use only a uniform bound. For any $p, q, r \in [0,1]$, letting $u = (p+q)/2$ and $v = (p+q+r)/3$ we have $u \geq p/2$, $1-u \geq (1-p)/2$ and $v \geq p/3$, $1-v \geq (1-p)/3$, hence

$$\mathrm{KL}(\mathrm{Ber}(p)\|\mathrm{Ber}(u)) \leq \log 2, \qquad \mathrm{KL}(\mathrm{Ber}(p)\|\mathrm{Ber}(v)) \leq \log 3.$$

Therefore $\mathcal{D}_{32} \leq 2\log 2$ and $\mathcal{D}_{31} \leq 3\log 3$ always, and thus

$$\mathbb{E}[\mathcal{D}_{32}\mathbf{1}_B \mid x] \leq 2\log 2\,\mathbb{P}(B \mid x), \qquad \mathbb{E}[\mathcal{D}_{31}\mathbf{1}_B \mid x] \leq 3\log 3\,\mathbb{P}(B \mid x).$$

It remains to show $\mathbb{P}(B \mid x) = o(F_{root} + F_{sub} + F_{out})$. This probability is exponentially small, because Lemma 7 applied to $P_1^0 \mid x$ and $P_{23}^0 \mid x$ gives $\mathbb{P}(P_1^0 \notin [\varepsilon_0, 1-\varepsilon_0] \mid x) \leq Ce^{-c/F_{out}}$ and $\mathbb{P}(P_{23}^0 \notin [\varepsilon_0, 1-\varepsilon_0] \mid x) \leq Ce^{-c/F_{root}}$. For $P_2^0$ and $P_3^0$, fix $\varepsilon_1 \in (\varepsilon_0, \varepsilon)$ (e.g. $\varepsilon_1 = (\varepsilon+\varepsilon_0)/2$). Then

$$\mathbb{P}(P_2^0 \notin [\varepsilon_0, 1-\varepsilon_0] \mid x) \leq \mathbb{P}(P_{23}^0 \notin [\varepsilon_1, 1-\varepsilon_1] \mid x) + \sup_{y \in [\varepsilon_1, 1-\varepsilon_1]} \mathbb{P}(P_2^0 \notin [\varepsilon_0, 1-\varepsilon_0] \mid P_{23}^0 = y), \quad (62)$$

which follows by splitting on the event $\{P_{23}^0 \in [\varepsilon_1, 1-\varepsilon_1]\}$. Indeed, letting $A = \{P_2^0 \notin [\varepsilon_0, 1-\varepsilon_0]\}$ and $C = \{P_{23}^0 \in [\varepsilon_1, 1-\varepsilon_1]\}$,

$$\mathbb{P}(A \mid x) = \mathbb{P}(A \cap C \mid x) + \mathbb{P}(A \cap C^c \mid x) \leq \mathbb{P}(C^c \mid x) + \mathbb{P}(A \cap C \mid x),$$

and

$$\mathbb{P}(A \cap C \mid x) \leq \sup_{y \in [\varepsilon_1, 1-\varepsilon_1]} \mathbb{P}(A \mid P_{23}^0 = y).$$

Lemma 7 (first with $(\varepsilon_1, \varepsilon)$ and then with $(\varepsilon_0, \varepsilon_1)$) bounds the right-hand side of (62) by $Ce^{-c/F_{root}} + Ce^{-c/F_{sub}}$; the same holds for $P_3^0$. Hence $\mathbb{P}(B \mid x) \leq C\exp(-c/\max\{F_{root}, F_{sub}, F_{out}\})$ by a union bound, and so $\mathbb{P}(B \mid x) = o(F_{root} + F_{sub} + F_{out})$ as $(F_{root}, F_{sub}, F_{out}) \to 0$. In particular,

$$\mathbb{E}[\mathcal{D}_{32} \mid x] = \mathbb{E}[\mathcal{D}_{32}\mathbf{1}_G \mid x] + o(F_{root} + F_{sub} + F_{out}),$$

and

$$\mathbb{E}[\mathcal{D}_{31} \mid x] = \mathbb{E}[\mathcal{D}_{31}\mathbf{1}_G \mid x] + o(F_{root} + F_{sub} + F_{out}),$$

so it suffices to analyze the expectations on $G$.

Recall that the Balding-Nichols model is a particular way of parameterizing the Beta distribution so that $\mathbb{E}[P_1^0|x] = \mathbb{E}[P_{23}^0|x] = x$,

$$\mathrm{Var}(P_1^0|x) = F_{out}x(1-x), \quad \mathrm{Var}(P_{23}^0|x) = F_{root}x(1-x),$$

and

$$\mathrm{Var}(P_2^0|P_{23}^0) = \mathrm{Var}(P_3^0|P_{23}^0) = F_{sub}P_{23}^0(1-P_{23}^0).$$

**Step 1: leading term for $\mathcal{D}_{32}$.**

On $G$, we have $\varepsilon_0 \leq P_2^0, P_3^0, m_{23} \leq 1-\varepsilon_0$. Thus Lemma 6 (applied with $\varepsilon_0$) to $(P_2^0, m_{23})$ and $(P_3^0, m_{23})$ yields

$$\begin{aligned}\mathrm{KL}(\mathrm{Ber}(P_2^0)\|\mathrm{Ber}(m_{23})) &+ \mathrm{KL}(\mathrm{Ber}(P_3^0)\|\mathrm{Ber}(m_{23})) \\ &= \frac{(P_2^0 - m_{23})^2 + (P_3^0 - m_{23})^2}{2m_{23}(1-m_{23})} + O_{\varepsilon_0}(|P_2^0 - P_3^0|^3). \quad (63)\end{aligned}$$

Since $P_2^0 - m_{23} = (P_2^0 - P_3^0)/2$ and $P_3^0 - m_{23} = -(P_2^0 - P_3^0)/2$, the numerator equals $(P_2^0 - P_3^0)^2/2$. Moreover, conditional on $x$, we have $\mathbb{E}[m_{23} \mid x] = x$ and

$$\mathrm{Var}(m_{23} \mid x) = \mathrm{Var}(P_{23}^0 \mid x) + \mathbb{E}[\mathrm{Var}(m_{23} \mid P_{23}^0) \mid x] = F_{root}x(1-x) + \frac{1}{2}F_{sub}(1-F_{root})x(1-x),$$

so $m_{23} - x = O_p(\sqrt{F_{root} + F_{sub}})$. Since $g(t) = 1/(t(1-t))$ has bounded derivative on $[\varepsilon_0, 1-\varepsilon_0]$, we also have

$$\frac{1}{m_{23}(1-m_{23})} = \frac{1}{x(1-x)} + O_p(|m_{23} - x|) = \frac{1}{x(1-x)} + O_p(\sqrt{F_{root} + F_{sub}}). \tag{64}$$

Next, we claim that

$$\mathbb{E}[(P_2^0 - P_3^0)^4 \mid x] = O(F_{sub}^2). \tag{65}$$

Indeed, conditional on $P_{23}^0 = y$, the variables $P_2^0$ and $P_3^0$ are i.i.d. $\mathrm{Beta}(\alpha_{sub}y, \alpha_{sub}(1-y))$ with $\alpha_{sub} = (1-F_{sub})/F_{sub}$ and mean $y$, so writing $U = P_2^0 - y$ and $V = P_3^0 - y$ gives

$$\begin{aligned}
\mathbb{E}[(P_2^0 - P_3^0)^4 \mid y] &= \mathbb{E}[(U-V)^4 \mid y] \\
&= \mathbb{E}[U^4 \mid y] - 4\mathbb{E}[U^3 V \mid y] + 6\mathbb{E}[U^2V^2 \mid y] - 4\mathbb{E}[UV^3 \mid y] + \mathbb{E}[V^4 \mid y] \\
&= 2\mathbb{E}[U^4 \mid y] + 6\,\mathbb{E}[U^2 \mid y]^2.
\end{aligned}$$

Here we used that $U$ and $V$ are i.i.d. and independent given $y$ with $\mathbb{E}[U \mid y] = \mathbb{E}[V \mid y] = 0$, so $\mathbb{E}[U^3V \mid y] = \mathbb{E}[U^3 \mid y]\mathbb{E}[V \mid y] = 0$, $\mathbb{E}[UV^3 \mid y] = 0$, and $\mathbb{E}[U^2V^2 \mid y] = \mathbb{E}[U^2 \mid y]\mathbb{E}[V^2 \mid y] = \mathbb{E}[U^2 \mid y]^2$. Moreover, $\mathbb{E}[U^2 \mid y] = \mathrm{Var}(P_2^0 \mid y) = F_{sub}y(1-y)$ and the Beta kurtosis is uniformly bounded when $y \in [\varepsilon_0, 1-\varepsilon_0]$, implying $\mathbb{E}[U^4 \mid y] = O(\mathrm{Var}(P_2^0 \mid y)^2) = O(F_{sub}^2)$. Averaging over $y$ yields $\mathbb{E}[(P_2^0 - P_3^0)^4 \mid x] = O(F_{sub}^2)$.

Hence, multiplying the error term by $(P_2^0 - P_3^0)^2$, which is of size $O(F_{sub})$ in expectation, we get by Cauchy–Schwarz,

$$\begin{aligned}
\mathbb{E}\big[(P_2^0 - P_3^0)^2|m_{23} - x| \mid x\big] &\le \mathbb{E}[(P_2^0 - P_3^0)^4 \mid x]^{1/2}\,\mathbb{E}[(m_{23} - x)^2 \mid x]^{1/2} \\
&= O(F_{sub})\,O(\sqrt{F_{root} + F_{sub}}) = o(F_{root} + F_{sub}).
\end{aligned}$$

Combine with (64), it implies that replacing $m_{23}(1-m_{23})$ by $x(1-x)$ in (63) changes the expectation only by $o(F_{root} + F_{sub})$. Moreover, the cubic remainder term is negligible because of Cauchy–Schwarz:

$$\mathbb{E}\big[|P_2^0 - P_3^0|^3 \mid x\big] \le \mathbb{E}\big[(P_2^0 - P_3^0)^4 \mid x\big]^{1/2}\,\mathbb{E}\big[(P_2^0 - P_3^0)^2 \mid x\big]^{1/2} = O\Big(F_{sub}^{3/2}\Big),$$

so the contribution of $O_{\varepsilon_0}(|P_2^0 - P_3^0|^3)$ is $o(F_{root} + F_{sub})$ as $(F_{root}, F_{sub}) \to 0$. Finally,

$$\mathbb{E}\big[(P_2^0 - P_3^0)^2 \mid x\big] = 2\,\mathbb{E}\big[\mathrm{Var}(P_2^0 \mid P_{23}^0) \mid x\big] = 2F_{sub}\,\mathbb{E}[P_{23}^0(1-P_{23}^0) \mid x] = 2F_{sub}(1-F_{root})x(1-x),$$

where the last identity uses $\mathbb{E}[P_{23}^0(1-P_{23}^0) \mid x] = x(1-x) - \mathrm{Var}(P_{23}^0|x) = (1-F_{root})x(1-x)$. This yields (57) after averaging over loci and $P^*$ and collecting remainder terms.

**Step 2: leading term for $\mathcal{D}_{31}$.**

On $E_{\varepsilon_0}$ we have $\varepsilon_0 \le m \le 1-\varepsilon_0$, so Lemma 6 (with $\varepsilon_0$) applied to each $(P_k^0, m)$ gives

$$\sum_{k=1}^{3} \mathrm{KL}(\mathrm{Ber}(P_k^0)\|\mathrm{Ber}(m)) = \frac{\sum_{k=1}^{3}(P_k^0 - m)^2}{2m(1-m)} + O_{\varepsilon_0}\left(\sum_{k=1}^{3}|P_k^0 - m|^3\right).$$

Using the identity

$$\sum_{k=1}^{3}(P_k^0 - m)^2 = \frac{2}{3}(P_1^0 - m_{23})^2 + \frac{1}{2}(P_2^0 - P_3^0)^2,$$

it remains to compute the conditional second moments. Since $P_1^0$ is independent of $(P_2^0, P_3^0)$ given $x$, we have

$$\mathbb{E}[(P_1^0 - m_{23})^2 \mid x] = \mathrm{Var}(P_1^0|x) + \mathrm{Var}(m_{23} \mid x).$$

By the law of total variance, $\mathrm{Var}(m_{23} \mid x) = \mathrm{Var}(P_{23}^0 \mid x) + \mathbb{E}[\mathrm{Var}(m_{23} \mid P_{23}^0) \mid x]$, and

$$\mathrm{Var}(m_{23} \mid P_{23}^0) = \mathrm{Var}\left(\frac{P_2^0 + P_3^0}{2} \;\middle|\; P_{23}^0\right) = \frac{1}{2}F_{sub}P_{23}^0(1 - P_{23}^0).$$

Therefore,

$$\mathrm{Var}(m_{23} \mid x) = F_{root}x(1-x) + \frac{1}{2}F_{sub}(1 - F_{root})x(1-x).$$

Combining with $\mathrm{Var}(P_1^0|x) = F_{out}x(1-x)$ and the expression for $\mathbb{E}[(P_2^0 - P_3^0)^2 \mid x]$ computed in Step 1 yields

$$\mathbb{E}\left[\sum_{k=1}^{3}(P_k^0 - m)^2 \;\middle|\; x\right] = \left(\frac{2}{3}(F_{out} + F_{root}) + \frac{4}{3}F_{sub}(1 - F_{root})\right)x(1-x) + o(F_{root} + F_{sub} + F_{out}).$$

Replacing $m(1-m)$ by $x(1-x)$ in the denominator (again contributing only higher-order terms in expectation) and averaging over loci gives (58). Finally, substituting $F_{out} = F_{sub} + F_{root} - F_{sub}F_{root}$ gives (59).

### 3 A simple result on bounding KL divergence by $F_2$ and $F_3$ statistics

In this section, we examine how the asymptotic lower bound of $\widehat{\mathcal{L}}^{(3)} - \widehat{\mathcal{L}}^{(1)}$:

$$\mathcal{D}_{31} := \overline{\mathrm{KL}}\left(\mathbf{P}_1^0 \| \tfrac{\mathbf{P}_1^0 + \mathbf{P}_2^0 + \mathbf{P}_3^0}{3}\right) + \overline{\mathrm{KL}}\left(\mathbf{P}_2^0 \| \tfrac{\mathbf{P}_1^0 + \mathbf{P}_2^0 + \mathbf{P}_3^0}{3}\right) + \overline{\mathrm{KL}}\left(\mathbf{P}_3^0 \| \tfrac{\mathbf{P}_1^0 + \mathbf{P}_2^0 + \mathbf{P}_3^0}{3}\right) \tag{66}$$

and the asymptotic upper bound of $\widehat{\mathcal{L}}^{(3)} - \widehat{\mathcal{L}}^{(2)}$:

$$\mathcal{D}_{32} := \overline{\mathrm{KL}}\left(\mathbf{P}_2^0 \| \tfrac{\mathbf{P}_2^0 + \mathbf{P}_3^0}{2}\right) + \overline{\mathrm{KL}}\left(\mathbf{P}_3^0 \| \tfrac{\mathbf{P}_2^0 + \mathbf{P}_3^0}{2}\right), \tag{67}$$

relate to distance notions frequently used in population genetics, such as $F_2$, and $F_3$ statistics. The $F_2$ statistics between $\mathbf{P}_1$ and $\mathbf{P}_2$ is denoted by

$$F_2(\mathbf{P}_1, \mathbf{P}_2) = \frac{1}{L}\sum_{\ell \in [L]}(P_{1\ell} - P_{2\ell})^2.$$

We can utilize the Pinsker's inequality to provide a lower bound for $\mathcal{D}_{31}$ in terms of $F_2$ statistics.

**Proposition 1.** *We have*

$$\sum_{k=1}^{3} \overline{KL}\left(\boldsymbol{P}_k^0 \| \tfrac{\boldsymbol{P}_1^0+\boldsymbol{P}_2^0+\boldsymbol{P}_3^0}{3}\right) \geq \frac{2}{3}\left(F_2(\boldsymbol{P}_1^0, \boldsymbol{P}_2^0) + F_2(\boldsymbol{P}_2^0, \boldsymbol{P}_3^0) + F_2(\boldsymbol{P}_1^0, \boldsymbol{P}_3^0)\right), \tag{68}$$

*or equivalently,*

$$\sum_{k=1}^{3} \overline{KL}\left(\boldsymbol{P}_k^0 \| \tfrac{\boldsymbol{P}_1^0+\boldsymbol{P}_2^0+\boldsymbol{P}_3^0}{3}\right) \geq \frac{4}{3}\left(F_3(\boldsymbol{P}_1^0; \boldsymbol{P}_2^0, \boldsymbol{P}_3^0) + F_3(\boldsymbol{P}_2^0; \boldsymbol{P}_1^0, \boldsymbol{P}_3^0) + F_3(\boldsymbol{P}_3^0; \boldsymbol{P}_1^0, \boldsymbol{P}_2^0)\right). \tag{69}$$

*Proof of Proposition 1.* Thanks to Pinsker's inequality, we have

$$\mathrm{KL}\left(\mathrm{Ber}(P_{k\ell}^0) \| \mathrm{Ber}\left(\tfrac{P_{1\ell}^0+P_{2\ell}^0+P_{3\ell}^0}{3}\right)\right) \geq 2\mathrm{V}^2\left(\mathrm{Ber}(P_{k\ell}^0), \mathrm{Ber}\left(\tfrac{P_{1\ell}^0+P_{2\ell}^0+P_{3\ell}^0}{3}\right)\right),$$

for every SNP $\ell \in [L]$ and population $k \in [3]$, where V is the Total Variation distance, and

$$\mathrm{V}^2\left(\mathrm{Ber}(P_{k\ell}^0), \mathrm{Ber}\left(\tfrac{P_{1\ell}^0+P_{2\ell}^0+P_{3\ell}^0}{3}\right)\right) = \left(P_{k\ell}^0 - \tfrac{P_{1\ell}^0+P_{2\ell}^0+P_{3\ell}^0}{3}\right)^2.$$

Therefore, by some simple algebraic manipulations,

$$\begin{aligned}\sum_{k=1}^{3} \mathrm{KL}\left(\mathrm{Ber}(P_{k\ell}^0) \| \mathrm{Ber}\left(\tfrac{P_{1\ell}^0+P_{2\ell}^0+P_{3\ell}^0}{3}\right)\right) &\geq 2\sum_{k=1}^{3}\left(P_{k\ell}^0 - \tfrac{P_{1\ell}^0+P_{2\ell}^0+P_{3\ell}^0}{3}\right)^2 \\ &= \frac{2}{3}\left((P_{1\ell}^0 - P_{2\ell}^0)^2 + (P_{2\ell}^0 - P_{3\ell}^0)^2 + (P_{3\ell}^0 - P_{1\ell}^0)^2\right)\end{aligned}$$

Taking average across all SNPs $\ell \in [L]$ we have (68). Moreover, notice that

$$F_3(\mathbf{P}_1^0; \mathbf{P}_2^0, \mathbf{P}_3^0) = \frac{1}{2}(F_2(\mathbf{P}_1^0, \mathbf{P}_2^0) + F_2(\mathbf{P}_1^0, \mathbf{P}_3^0) - F_2(\mathbf{P}_2^0, \mathbf{P}_3^0)). \tag{70}$$

We can write $F_3(\mathbf{P}_2^0; \mathbf{P}_1^0, \mathbf{P}_3^0)$ and $F_3(\mathbf{P}_3^0; \mathbf{P}_1^0, \mathbf{P}_2^0)$ similarly in terms of $F_2$ statistics. Hence,

$$F_3(\mathbf{P}_1^0; \mathbf{P}_2^0, \mathbf{P}_3^0) + F_3(\mathbf{P}_2^0; \mathbf{P}_1^0, \mathbf{P}_3^0) + F_3(\mathbf{P}_3^0; \mathbf{P}_1^0, \mathbf{P}_2^0) = \frac{1}{2}\left(F_2(\mathbf{P}_1^0, \mathbf{P}_2^0) + F_2(\mathbf{P}_1^0, \mathbf{P}_3^0) + F_2(\mathbf{P}_2^0, \mathbf{P}_3^0)\right). \tag{71}$$

Combining this equation with (68) we have (69). □

### 4 Discussion on related works and using $\Delta K$ with Bayesian methods

When making inference of the admixture model using the Bayesian method [Pritchard et al., 2000] (implemented in the program STRUCTURE) for $K$ populations, we obtain the posterior sampling

$$\{(Z^{(m)}, P^{(m)}, Q^{(m)}) : m \in [M]\}$$

of the latent variables $Z$ and model parameters $P, Q$, given observed genotype data $X$. The original paper [Pritchard et al., 2000] suggests approximating the model evidence $P(X|K)$ as follows. First we calculate the mean of the log-likelihood model with respect to the posterior samples:

$$\widehat{\mu}_K := \frac{1}{M}\sum_{m=1}^{M} \log p(X|Z^{(m)}, P^{(m)}, Q^{(m)}), \tag{72}$$

and its variance:

$$\widehat{\sigma}_K^2 := \frac{1}{M} \sum_{m=1}^{M} \left( \log p(X|Z^{(m)}, P^{(m)}, Q^{(m)}) - \widehat{\mu}_K \right)^2 \tag{73}$$

The log evidence is approximated by:

$$\log P(X|K) \approx \widehat{\mu}_K - \frac{\widehat{\sigma}_K^2}{2} =: L_K. \tag{74}$$

The second-order of change for $L_K$ is defined as

$$L_K'' = |2L_K - L_{K-1} - L_{K+1}|. \tag{75}$$

This procedure is then replicated $B$ times (say, $B = 20$ times of running MCMC) to obtain a set of values of $L_K$. Let $m(L_K'')$ denote the mean of $L_K''$ in those $B$ runs and $s(L_K)$ denote the standard deviation of $L_K$. The $\Delta K$ is defined as

$$\Delta K := m(L_K'')/s(L_K). \tag{76}$$

In this paper we study the limit of $L_K''$ as if $L_K$ is defined by the log-likelihood of the MLE $L_K = \log p(X|\widehat{P}^{(K)}, \widehat{Q}^{(K)})$. This is a natural adaptation of Evanno's method for MLE, since we do not have replications and posterior variance when inferring parameters in the admixture model with MLE. Our results shed light on the nested population structure case where selecting models by the second-order change of log-likelihood of the MLE might be inconsistent.

In the Bayesian formula, the limit of the log-likelihood (72) could be studied, for example, by using Bernstein von-Mises theorem [Ghosal and Van der Vaart, 2017], which will imply the limit of the second-order change $L_K''$. But it is challenging to analyze the limiting behavior of the standard deviation $s(L_K)$. Although it is slightly different from the original method proposed by [Evanno et al., 2005], the model selection method adapted in this paper shares the same intuition: As we increase $K$, the models' log-likelihood increases, and the increment (first-order change) from under-fitted level to exact-fitted level (1 to $K_0$) is different from that from exact-fitted level to exact-fitted level ($K_0$ to $\overline{K}$), therefore taking the second-order change can help detect the true $K_0$. We conduct a theoretical analysis of the MLE to demonstrate that this intuition does not consistently select the correct $K_0$ in the presence of nested population structure, where the increment in models' log-likelihood is not uniform from 1 to $K_0$.

### 5 Technical lemmas

In the proofs of Lemma 2, 4, and 5, we use many forms of Chernoff and Hoeffding's inequality for a Bernoulli sequence of random variables. They can be found in [Mitzenmacher and Upfal, 2017] (Chapter 4.2) or in [Vershynin, 2018]. We recall them here.

**Lemma 8.** *Suppose that $Y_i \sim \text{Ber}(p_i), i = 1, \ldots, N$ are independent. Let $\overline{Y} = \sum Y_i/N$ and $\overline{\mu} = \sum p_i/N$. We have the following concentration inequalities.*

1. *(Chernoff's inequality for relative error) For all $t > 0$:*

$$\mathbb{P}\left(|\overline{Y} - \overline{\mu}| \geq t\overline{\mu}\right) \leq 2e^{-N\overline{\mu}((t+1)\log(t+1)-t)}. \tag{77}$$

*Specifically, for any $\delta \in (0, 1)$*

$$\mathbb{P}\left(|\overline{Y} - \overline{\mu}| \geq \delta\overline{\mu}\right) \leq 2e^{-N\overline{\mu}\delta^2/3}. \tag{78}$$

2. *(Hoeffding's inequality) For any $t > 0$:*

$$\mathbb{P}\left(|\overline{Y} - \overline{\mu}| \geq t\right) \leq e^{-2Nt^2}. \tag{79}$$

**Lemma 9** (Bernstein's inequality, Theorem 2.8.4 [Vershynin, 2018])**.** *Let $Y_1, \ldots, Y_N$ be independent, mean zero random variables, such that $|Y_i| \leq M$ for all $i$. Then, for all $t \geq 0$,*

$$\mathcal{P}\left(\left|\sum_{i=1}^{N} Y_i\right| \geq t\right) \leq 2\exp\left(\frac{-t^2}{2(\sigma^2 + Mt/3)}\right), \tag{80}$$

*where $\sigma^2 = \sum_{i=1}^{N} \mathbb{E}Y_i^2$ is the variance of the sum.*
